# Summary-data-based mendelian randomisation reveals druggable targets for multiple sclerosis

**DOI:** 10.1101/2020.01.20.907451

**Authors:** Benjamin Meir Jacobs, Thomas Taylor, Amine Awad, David Baker, Gavin Giovanonni, Alastair Noyce, Ruth Dobson

**Author notes:** equal contribution (Thomas Taylor, Amine Awad, Alastair Noyce, Ruth Dobson).

## Abstract

**Background:** Multiple Sclerosis (MS) is a complex autoimmune disease caused by a combination of genetic and environmental factors. Translation of Genome-Wide Association Study (GWAS) findings in MS into therapeutics and effective preventive strategies has been limited to date.

**Methods:** We used Summary Data-Based Mendelian Randomisation (SMR) to synthesise findings from public expression quantitative trait locus (eQTL; eQTLgen and CAGE), methylation quantitative trait locus (mQTL; Lothian Birth Cohort and Brisbane Systems Genetics Study), and MS GWAS datasets (International Multiple Sclerosis Genetics Consortium). By correlating the effects of methylation on MS (M-2-MS), methylation on expression (M-2-E), and expression on MS susceptibility (E-2-MS), we prioritise genetic loci with strong evidence of causally influencing MS susceptibility. We overlay these findings onto a list of ‘druggable’ genes, i.e. genes which are currently, or could theoretically, be targeted by therapeutic compounds. We use GeNets and STRING to identify protein-protein interactions and druggable pathways enriched in our results. We extend these findings to a model of Epstein-Barr Virus-infected B cells, Lymphoblastoid Cell Lines (LCLs). We conducted a systematic review of prioritised genes using the Open Targets platform to identify completed and planned trials targeted prioritised genes in MS and related disease areas.

**Results:** Expression of 45 genes in peripheral was strongly associated with MS susceptibility (False discovery rate 0.05). Of these 45 genes, 20 encode a protein which is currently targeted by an existing therapeutic compound. These genes were enriched for Gene Ontology terms pertaining to immune system function and leukocyte signalling. We refined this prioritised gene list by restricting to loci where CpG site methylation was associated with MS susceptibility (M-2-MS), with gene expression (M-2-E), and where expression was associated with MS susceptibility (E-2-MS). This approach yielded a list of 15 prioritised druggable target genes for which there was evidence of a causal pathway linking methylation, expression, and MS. Five of these 15 genes are targeted by existing drugs (CD40, ERBB2, VEGFB, MERTK, and PARP1), and three were replicated in a smaller eQTL dataset (CD40, MERTK, and PARP1). In LCLs, SMR prioritised 7 druggable gene targets, of which only one was priortised by the multi-omic approach in peripheral blood (FCRL3). Systematic review of Open Targets revealed multiple early-phase trials targeting 13/20 prioritised genes in disorders related to MS.

**Conclusions:** We use public datasets and SMR to identify a list of prioritised druggable genetic targets in Multiple Sclerosis. We hope our findings could be translated into effective repurposing of existing drugs to provide novel therapies for MS and, potentially, provide a platform for developing preventive therapies.

## Introduction

Genome-wide association studies (GWAS) in Multiple Sclerosis (MS) have revealed over 200 risk loci associated with an increased risk of developing the disease^1^. However, translating findings from GWAS into therapeutic strategies has proved challenging for a number of reasons. GWAS provide insights into potential genomic risk loci that are likely to harbour single or multiple causal variants. Despite the advent of analytical techniques such as fine-mapping, advanced annotation tools, colocalisation, and mendelian randomisation for inferring causal variants from risk loci, difficulties remain in inferring with certainty which variants are truly causal. Understanding how these variants mechanistically influence disease phenotypes provides additional challenges^2^. These challenges arise from factors such as complex linkage disequilibrium and potential effects on distant (i.e. trans) genes. Additionally, dynamic, context-specific effect of variants are likely to vary depending on time, cell type, and context.

MS is a complex autoimmune disease, which is a leading cause of disability in young people. There are currently neither effective cures nor preventive measures for MS. Licensed therapies for MS include rationally-designed monoclonal antibodies, such as ocrelizumab and ofatumumab (anti-CD20), natalizumab (anti-alpha4 integrin) and alemtuzumab (anti-CD52), and fingolimod (sphingosine-1-phosphate-receptor modulators), in addition to drugs with multiple mechanisms of action such as cladribine, glatiramer acetate, and beta-interferon^3^. Autologous haematopoietic stem cell transplant (aHSCT) has in increasing evidence base for use in MS. Existing highly-active therapies are effective at controlling inflammatory disease activity, but have several serious side-effects; alongside this, prognostication at the time of diagnosis remains imperfect. There is thus an unmet clinical need for highly-effective therapies with a better safety profile, which is even more important when considering preventive drugs for individuals at high-risk of disease.

Many GWAS hits reside in intronic or intergenic regions, or within genes that do not make attractive druggable targets, for example due to conformational considerations, cellular localisation, or concerns about off-target effects. Outside of rare, missense or nonsense coding variants, moving from GWAS hits into druggable targets has had limited success. Expression Quantitative Trait Loci (eQTL) studies provide a tissue-specific measure of genetic expression and enable a deeper understanding of the influence of transcription levels alongside genetic variation. Summary-data based mendelian randomization (SMR) provides a data-driven approach for integrating GWAS data with gene expression data from expression quantitative trait locus (eQTL) studies to prioritise potentially causal genes from GWAS hits^4^. SMR extends the concept of mendelian randomisation, enabling testing of the hypothesis that genetically-determined levels of gene expression are associated with a disease phenotype.

In this paper, we focus on a previously curated list of 4479 genes that were identified as ‘druggable’ on the basis of various considerations^5^. We apply SMR to recent MS GWAS data, complemented by pathway analysis to provide a data-driven prioritisation of druggable genes in MS.

## Methods

### Datasets

MS GWAS data were provided by the International MS Genetics Consortium (IMSGC) from the latest IMSGC discovery-stage summary statistics^1^. These data originate from 14,802 individuals with MS and 26,703 controls of European descent. Allele frequencies for SNPs in the MS GWAS discovery data were obtained from the 1000 genomes samples of European ancestry (n=503)^6^.

For SMR analyses, we used several expression Quantitative Trait Loci (eQTL) datasets. The primary analysis used cis-eQTL data from the eQTLgen consortium. This dataset contains cis-eQTLs for all 19,250 genes expressed in whole blood obtained from 31,684 individuals. Each SNP-gene pair for which data was available in ≥2 cohorts and with a SNP-gene distance of ≤ 1MB was tested^7^. Data were downloaded from the eQTLgen consortium website (https://www.eqtlgen.org/cis-eqtls.html)^7^. Whole blood results obtained with the eQTLgen dataset were replicated using the Consortium for the Architecture of Gene Expression (CAGE) consortium eQTL data obtained from peripheral blood of 2,765 individuals ^8^. CAGE data were downloaded from the SMR website (https://cnsgenomics.com/software/smr/#DataResource). Data on eQTLs in Lymphoblastocytoid Cell Lines (LCLs) - EBV-immortalised B cells - were obtained from the Geuvadis consortium results available from the SMR website ^9^.

For multi-omic SMR, we used peripheral blood methylation Quantitative Trait Locus (mQTL) data from a meta-analysis of the Lothian Birth Cohort (LBC) and the Brisbane Systems Genetics Study (BSGS)^2,10^. These data are limited to DNA methylation probes with ≥1 cis-mQTL associated at P < 5×10^−8^ and SNPs ≤ 2MB from each DNA methylation probe. mQTL data were downloaded from the SMR website.

### Druggable genome

The druggable genome can be defined as the set of protein-coding genes for which the gene products could potentially be modulated by therapeutic compounds. This includes protein products which are already targeted by existing drugs and proteins with structural and functional properties suggestive of druggability but which are not currently targeted by existing compounds. The list of druggable genes used in this work was taken from table S1 of the paper by Finan *et al*^5^. The final list was developed from a list of protein-coding genes, T cell receptor genes, immunoglobulins, polymorphic pseudogenes, and selected non-protein-coding genes believed to have functional consequences. Genes were classified into three tiers based on their druggability. Genes were classified as ‘Tier 1’ if they were already being targeted by compounds in clinical use or clinical development. ‘Tier 2’ genes were not currently targeted by existing compounds but have a peptide sequence product with high sequence homology to ‘Tier 1’ druggable genes. ‘Tier 3’ genes incorporated gene products with a degree of peptide sequence homology to targets of existing compounds, genes encoding major classes of druggable protein (kinases, ion channels, G-protein coupled receptors, nuclear hormone receptors, and phosphodiesterases), genes encoding extracellular proteins (either secreted or membrane-bound), and Cluster of Differentiation (CD) antigen genes. Tier 3 was divided into 3A and 3B based on proximity to GWAS hits for various common diseases, with genes <=50KB from a GWAS hit deemed more likely to be druggable (3A).

### SMR

Summary-data based mendelian randomisation (SMR) is a technique used to determine associations between genetically-determined traits, such as gene expression and methylation, and outcomes of interest, such as disease phenotypes^4^. eQTLs refer to genetic variants which are associated with levels of expression of a particular transcript. These are derived from the measurement of gene expression, most commonly using RNA sequencing, which is then correlated with genotyping data. Importantly, eQTLs are time and tissue-specific: the genetic regulation of gene expression varies widely between tissues and depends on context.

In SMR, SNP association statistics with an outcome (e.g. a disease phenotype) are regressed on SNP association statistics with expression of a particular transcript to determine a ‘causal estimate’, approximating the effect on the disease phenotype for a genetically-determined increase in the expression of that transcript. If *β*_E_ is the per-allele beta for a genetically-determined increase in gene expression, and *β*_D_ is the per-allele log odds ratio for a binary disease phenotype, then the causal estimate for the effect of genetically-determined increased expression of that transcript *β*_SMR_ is *β*_D_ / *β*_E_. A key assumption of this approach is that the same underlying causal variant determines both gene expression and the disease phenotype. Due to linkage disequilibrium, it is possible that *β*_SMR_ could be non-zero even when this assumption is violated, i.e if SNP *i* is in linkage with SNP *j*, which determines the expression of transcript *t*, and is also in linkage with SNP *k*, which directly influences the disease phenotype, then *β*_SMR_ may be non-zero despite no direct causal pathway from SNP to transcript to disease (see figure 1 in Wu *et al* ^2^). This is different to the ‘vertical pleiotropy’ situation upon which instrumental variable analysis and MR are based, which assumes a direct causal pathway between genetic variant, gene expression, and disease phenotype.

To distinguish vertical pleiotropy from linkage, Zhu *et al* developed the heterogeneity in dependent instruments (HEIDI) test, which exploits the observation that if gene expression and disease phenotype are in vertical pleiotropy with the same causal variant, *β*_SMR_ is identical for any variant in LD with the causal variant^4^. Thus greater heterogeneity among *β*_SMR_ statistics calculated for all significant cis-eQTLs implies a greater likelihood that linkage, rather than causality/vertical pleiotropy, explains the observed *β*_SMR_. The heterogeneity statistic, the ‘HEIDI’ statistic, tests the hypothesis HEIDI 0. This provides a formal test of heterogeneity, with *p* values < 0.05 suggestive of linkage, rather than vertical pleiotropy and a causal pathway^4^.

In this work we performed SMR using the SMR software tool (SMR v1.0.2) in the command line using default options^4^. Default options are as follows: cis-eQTLs selected based on minimum *p*=5×10^−8^, eQTLs included for the HEIDI test based on minimum *p*=1.57×10^−3^, eQTLs included for the HEIDI test if R^2^ with the top cis-eQTL was between 0.05 and 0.9, minimum number of SNPs included in the HEIDI test = 3, maximum number of SNPs included in the HEIDI test = 20, and physical window around probe within which the top cis-eQTL was selected = 2 MB.

P values were adjusted in R (v3.6.1) to control the false discovery rate at LJ = 0.05. Associations with *p_HEIDI_*<0.01 - the cutoff used by Wu *et al*^2^-were considered likely due to linkage and thus discarded from the analysis. Probes were excluded if any of the transcript or the top eQTL resided within the super-extended Major Histocompatibility Complex (MHC; hg19 6:25,000,000 - 35,000,000) given the complex LD structures within this region. LD estimation was performed using reference genomes obtained from the 1000 genomes samples of European ancestry (n=503)^6^.

### Multi-omic SMR

To further refine the list of plausible druggable targets developed using the techniques above, we performed multi-omic SMR using the methods described in Wu *et al* ^2^. This approach prioritises genes by layering SNP associations with CpG methylation sites, gene expression, and the phenotype of interest. As the majority of GWAS hits are in non-coding regions, they are likely to influence disease through regulation of gene expression^2^. As CpG methylation sites are partly genetically-determined, methylome-wide association study (MWAS) data can be exploited to determine likely causal relationships between CpG methylation and other traits, including molecular traits such as gene expression. We applied the following steps to prioritise druggable genes in MS:

1. Use SMR to determine associations between CpG methylation sites and gene expression (M → E)
2. Use SMR to determine associations between CpG methylation sites and MS (M → D)
3. Use SMR to determine associations between gene expression and MS (E → D)

If there is strong evidence of causal association (i.e. *p*_SMR_ < threshold and *p*_HEIDI_ > 0.01) at each of the above 3 steps (M → E, M → D, and E → D), this provides evidence of a causal pathway linking genetic variation, methylation, gene expression, and MS at that genetic locus.

Custom gene tracks for locus plots were downloaded from the USCS Genome Browser (https://genome.ucsc.edu/cgi-bin/hgTables). All genomic co-ordinates specified are genome build hg19 (GRCh37).

### Pathway analysis, functional annotation, and prediction of protein interactors

Pathway analysis and annotation was conducted using the STRING database of protein-protein interactions (https://string-db.org/). For data visualisation, we used default settings to map protein-protein interactions between the prioritised druggable targets. STRING database interaction scores are curated from multiple sources (experimental data, co-expression, text-mining, and predictions from peptide sequences) and reflect the probability that two proteins are linked in the same KEGG metabolic pathway. By default, interactions with medium confidence (>0.4) are shown^11,12^. For functional enrichment analysis, we examined the list of SMR-prioritised genes (not restricted to the druggable genome).

Enrichment for Gene Ontology (GO) terms and biological pathways was determined by querying GO terms, KEGG pathways, and REACTOME pathways using STRING^13^. Enrichment P values are computed using hypergeometric tests and corrected using the Benjamini-Hochberg procedure. The hypergeometric test for enrichment is based on the observation that under the null hypothesis (no enrichment), the number of genes from the gene list which have a particular GO term attached follows an approximately binomial distribution for large n values^14^.

To identify additional druggable targets not prioritised in the primary analysis but likely to interact with prioritised genes, we used the GeNets tool (http://apps.broadinstitute.org/genets). GeNets uses pre-trained random forest classifiers to predict the likelihood that any given protein interacts functionally with another. In addition to calculating interaction strength between given gene products, GeNets can also predict likely interacting partners given a gene set. We ran GeNets using the online tool, with the SMR-prioritised gene list from our primary analysis (eQTLgen SMR, supplementary table 3) as input, and with default settings^15^.

### Systematic review of clinical trials

Finally, we performed a systematic evaluation of the clinical trial literature for all tier 1 druggable targets associated with MS from our primary analysis (eQTL SMR). We searched the Open Targets Platform (https://www.targetvalidation.org/) for each prioritised gene candidate. For each gene, we collated all trials targeting the gene in autoimmune diseases (e.g. MS, Rheumatoid Arthritis, Inflammatory Bowel Disease, Psoriasis, Ankylosing Spondylitis, Systemic Lupus Erythematosus, Primary Biliary Cholangitis, Type 1 Diabetes Mellitus, Sjogren’s Syndrome, and Primary Sclerosing Cholangitis) and haematological malignancies. These diseases areas were chosen given their potentially shared aetiology with MS; successful trials in these diseases may suggest possible therapeutic benefit in MS.

### Statistical analysis, software, and computing

All analyses were conducted using SMR and R v3.6.1. This research was supported by the High-Performance Cluster computing network hosted by Queen Mary University of London^16^. SMR locus plots were made using the online tool hosted at the SMR website (https://cnsgenomics.com/software/smr/omicsplot/).

## Results

### Expression of druggable gene targets is modulated by MS risk loci

We used SMR to test the association of genetically-determined expression of druggable genes in peripheral blood with multiple sclerosis. Expression of 45 tested probes (mapping to 45 unique genes; 2356 probes tested in total) was associated with MS (FDR_SMR_ 0.05, *p*_HEIDI_ > 0.01, figure 1). Twenty of these genes were Tier 1 druggable targets (figure 1, figure 2, table 1). 17 of these 45 signals were replicated in the CAGE dataset (supplementary tables 1 & 2, supplementary figure 1). For these replicated genes, the association statistics (beta_SMR_) with MS were highly correlated between the two datasets (Pearson’s correlation coefficient 0.93, 95% CI 0.83 - 0.98, p = 4.20×10^−8^, supplementary tables 1 & 2, supplementary figure 2).

We found evidence for multiple protein-protein interactions within this list of prioritised druggable genes (figure 3a). Extending the SMR approach beyond the druggable genome, expression of 235 genes was associated with MS risk (FDR_SMR_ < 0.05, p_HEIDI_ >0.01, supplementary table 3, figure 3b). This full set of SMR-prioritised genes was enriched for multiple GO terms and biological pathways implicated in immune system function and dysfunction (supplementary table 4 - 6). No significant enrichment was found for REACTOME pathways nor for GO cellular components.

### Methylation at CpG sites is modulated by MS risk loci

Next, in order to determine CpG methylation sites associated with MS risk, we applied SMR to CpG mQTL data from a meta-analysis of the LBC and BSGS cohorts (see methods). We found evidence for an association of 574 CpG methylation probes with MS risk (table 2).

### Multi-omic SMR to prioritise druggable gene candidates in MS

To determine the associations between CpG methylation sites and gene expression, we performed SMR using the LBC and BSGS mQTL meta-analysis and the eQTLgen consortium eQTL data. We restricted this analysis to druggable gene transcripts and excluded the MHC. We identified 23,731 associated pairs of CpG probes and transcripts (FDR_SMR_ < 0.05, *p*_HEIDI_ > 0.01) mapping to 2548 unique genes (supplementary table 7).

We then overlaid SMR evidence for causal relationships between methylation and expression (M → E), methylation and MS (M → D), and expression and MS (E → D) for the druggable genome. We found 35 druggable loci mapping to 15 unique genes (table 3). Five of these genes were Tier 1 (highly druggable) genes: *CD40*, *ERBB2*, *VEGFB*, *MERTK*, and *PARP1*. Eight of these 15 unique genes were replicated using the CAGE dataset, including three of the five Tier 1 genes (*CD40*, *MERTK*, and *PARP1*). In addition, another Tier 1 gene, *IL12RB1*, was associated with MS in the CAGE but not the eQTLgen dataset (table 3, supplementary table 8). Locus plots with chromatin state annotations from the Roadmap Epigenomics Projects are shown for the three replicated Tier 1 genes (figure 4).

### Prediction of novel interactions

To extend the list of functionally-prioritised druggable targets, we used GeNets to predict which proteins interact with the targets identified in our analysis. As input, we used the full list of 235 SMR-prioritised genes (including non-druggable targets) from the primary analysis (supplementary table 3). This approach yielded an additional 75 predicted protein interaction partners and multiple novel and established therapeutic compounds targeting our list of prioritised protein networks (supplementary figure 4, supplementary table 10).

### Druggable targets in lymphoblastoid cell lines (LCLs)

EBV-transformed cell lines, Lymphoblastoid Cell Lines (LCLs), provide a model for studying gene expression on B cells *in vitro*. These cells are immortalised using EBV, which apppears to play a significant role in MS risk. We attempted to identify druggable gene targets in LCLs using LCL-eQTL data from the Geuvadis consortium ^9^. SMR identified 7 prioritised druggable gene targets for MS in LCLs (supplementary table 10, supplementary figure 5). Expression of 3/7 of these transcripts (SLC12A7, FCRL3, and SLAMF7) was also associated with MS susceptibility in blood, with concordant directions of effects for all three genes. The FCRL3 gene was the only gene that overlapped with our list of genes prioritised using multi-omic SMR, although notably the TNFRSF14 gene prioritised from the LCL analysis is adjacent to MMEL1, which was nominated by the multi-omic method as a likely causal gene in peripheral blood. Comparison of SMR causal estimates for gene expression of the 1215 overlapping genes which passed the HEIDI test (p > 0.01) revealed a weak but precisely estimated positive correlation between association statistics with MS in blood and LCLs (Pearson’s rho = 0.27, 95% CI 0.21 - 0.32, p<2×10^−16^).

### Systematic review of clinical trials

We conducted a systematic review of clinical trials targeting our twenty Tier 1 prioritised genes from the primary analysis (eQTL SMR with eQTLgen data). Using the Open Targets platform, we retrieved trial data for trials in autoimmune diseases (including MS) and in haematological malignancies for each druggable target. Full results are presented in supplementary table 9. Of the twenty genes queried, we found drug trials in related disease areas for 13/20. Gene targets without any trials in relevant areas were MPO, ABCC2, CD5, MERTK, NCSTN, MAP3K11, and VEGFB. Only one gene target, S1PR1, was being currently targeted in MS. Phase III trial data have demonstrated efficacy of S1PR1 modulators (fingolimod, siponimod, ozanimod and posenimod) in reducing relapse rate for relapsing disease ^17–19^ and slowing disability progression for secondary-progressive disease^20^. Four drug targets had been targeted in trials for at least one autoimmune disease (excluding MS): IFNGR2, HDAC3, TYK2, and CD40. Of these targets, only compounds targeting TYK2 had progressed into phase III/IV trial development. Completed phase III trials have shown efficacy of TYK2 inhibitors (e.g. tofacitinib) in RA^21^, psoriasis/psoriatic arthropathy^22^, and Ulcerative Colitis^23^. The other genes prioritised and replicated by the multi-omic SMR approach - CD40 and PARP1 - were not being targeted by any compounds beyond phase II development. CD40 inhibitors (lucatumumab and dacetuzumab) and PARP1 inhibitors (veliparib, niraparib, talazoparib) were in multiple phase I and II trials for haematological malignancies. An anti-CD40L mAb, INX-021,has entered Phase I testing in MS.

## Discussion

In this report we combine the latest IMSGC GWAS data with large mQTL and eQTL datasets and a list of curated ‘druggable genes’ to provide a data-driven list of 45 prioritised drug targets for multiple sclerosis. Three of these targets - CD40, MERTK, and PARP1 - have strong evidence to support a biological causal relationship between genetic variation, CpG site methylation, gene expression, and MS, and are already being targeted by existing compounds at various stages of clinical development. Several are being investigated in clinical or pre-clinical studies for multiple sclerosis and/or autoimmune disorders.

It is important to note that the phenotype examined in the GWAS used in this study is MS susceptibility (i.e. a binary disease trait), rather than MS severity. Thus, our findings should be interpreted as pointing to biological targets and pathways which could, in principle, be modulated to affect the risk of developing MS. While at least some of the pathways uncovered by this approach are also likely to play a role in determining MS severity, this assumption remains to be proven.

CD40 is a member of the Tumour Necrosis Factor (TNF) superfamily which is constitutively expressed on the cell surface of specialised antigen-presenting cells such as B cells and dendritic cells^24^. Binding of CD40 by its ligand, CD40L, initiates diverse signalling cascades via TNF Receptor-Associated-Factors (TRAFs) culminating in B cell differentiation, activation, proliferation, and germinal centre formation^24^. CD40L is primarily expressed on activated T cells, and the CD40L-CD40 interaction provides an antigen-specific costimulatory signal for propagating the humoral immune response. Several observations support a pathogenic role for CD40 signalling in MS: CD40L is upregulated in CD4^+^ T cells in MS lesions, soluble CD40L (a secreted ligand for CD40) is upregulated in serum and CSF of people with active MS, CD40-expressing mononuclear cells infiltrate the CNS in rodent experimental autoimmune encephalomyelitis (EAE), the density of CD40 appears higher on B cells from people with MS^24^, and variation at the *CD40* locus is associated with MS susceptibility^1^. The precise mechanism by which genetic variation at the *CD40* locus influences MS risk is unclear - perhaps counter-intuitively, MS-associated SNPs are associated with decreased expression of CD40 mRNA in peripheral blood and B lymphocyte subsets ^25,26^, an effect which is replicated in our study (Beta_SMR_ −0.50, p_SMR_ = 1.89×10^−12^). In a study of PBMCs from people with MS and healthy controls, the susceptibility SNP rs4810485 was associated with both alternative splicing of CD40 transcripts - with relative upregulation of the splice isoform lacking exon 5 leading to lower cell-surface expression - and with lower levels of the anti-inflammatory cytokine IL-10 in peripheral blood^26^. These data suggest that genetically-determined modulation of CD40 mRNA expression may promote MS risk via effects of alternatively-spliced transcripts on IL-10 production, although the exact mechanism remains to be clarified. Monoclonal antibodies blocking the interaction between CD40L and CD40 have shown efficacy in rodent and primate EAE^24^.

Drug trials of anti-CD40 therapies in humans have been limited by safety concerns. Early phase II data showed promising efficacy of an anti-CD40L mAb in lupus nephritis ^27^. Despite phase I safety of the anti-CD40L antibody, IDEC-131, in MS, the phase II trial was halted due to increased risk of thromboembolic disease caused by inhibition of the thrombus-stabilising interaction between CD40L and platelet alpha2,beta3 integrins^24,28^. Subsequently, various strategies have been deployed to overcome this safety concern. The thrombogenicity of CD40L antagonists is dependent on an Fc-FcγRIIa interaction leading to platelet cross-linking^29^; development of a PEGylated monovalent anti-CD40L Fab fragment lacking an Fc portion appears to have overcome this problem^30^. A placebo-controlled phase II trial of dapirolizumab pegol for individuals with SLE has been completed (results pending; NCT 02804763). An anti-CD40L antibody-like molecule based on a Tn3 protein scaffold has shown phase I safety and preliminary efficacy in individuals with rheumatoid arthritis^31^.

In addition to directly targeting the CD40L-CD40 interaction, it may be beneficial to target downstream signalling. Upon activation by CD40L, CD40 signals via nuclear factor kappa-light-chain-enhancer of activated B cells (NFkB), Phosphoinositide 3-kinases (PI3K), and the mitogen-activated protein kinases (MAPKs) c-Jun N-terminal kinase (JNK) and p38 MAPKs, depending on cellular context^32^. The potential utility of drugging the MAPK signalling pathway is clear from our study, with MAPK3, MAPK1, and MAP3K11 all prioritised in the primary analysis. Phosphoproteomic scans of differences in protein phosphorylation in pwMS have revealed constitutive high levels of activation of MAPK signalling in immune cells from pwMS^33^. Inhibitors of MEK1, a downstream target of MAPK signalling, have been trialled in rheumatoid arthritis, Crohn’s disease, psoriasis, and multiple myeloma^34^.

MERTK is a member of the TAM (Tyro3, Axl, and Mer) family of receptor tyrosine kinases which have multiple endogenous ligands, and exist in both membrane-bound and circulating (decoy receptor) forms^35^. Upon ligand binding, MERTK signals via PI3K/AKT, MAPK, and NFkB pathways, in addition to several others^35^. Physiological roles of MERTK include regulation of platelet function, inflammation, and phagocytosis^36^. Disruption of phagocytic clearance of dead and dying cells is thought to contribute to aberrant exposure of Damage-Associated Molecular Patterns (DAMPs), such as dsDNA, which may act as a substrate for loss of immune tolerance to self-antigens and subsequent autoimmunity^35^. In addition, MERTK appears to facilitate astrocytic clearance of synaptic debris in the CNS^37^. MERTK is a risk locus in the IMSGC GWAS^1^, with the MS risk-increasing alleles associated with increased MERTK expression (Beta_SMR_ 0.19, p_SMR_ 4.92×10^−8^). In cuprizone-treated rodents, knockout of the MERTK ligand, Growth Arrest Specific 6 (Gas6), delays remyelination and recovery, suggesting that the Gas6-MERTK interaction facilitates remyelination^38,39^. In post-mortem tissue, soluble and membrane-bound MERTK were up-regulated in chronic MS lesions; soluble MERTK was negatively correlated with lesional Gas6^40^. It is plausible that upregulation of MERTK leads to increased decoy receptor shedding (depending on metalloproteinase action) and subsequent inhibition of Gas6 by soluble MERTK. One study suggested that CSF Gas6 levels may be elevated in early relapse, but do not differ between controls and pwMS outside of an acute relapse^41^, which may reflect a homeostatic response to acute demyelination. Small molecule inhibitors of MERTK have shown efficacy in mouse models of ALL^42^.

Poly(ADP-ribose) polymerase 1*(PARP1)* is an intracellular enzyme which catalyses the transfer of ADP-ribose groups from NAD to targets such as histones, DNA repair proteins, and transcription factors. PARP1 is involved in DNA repair and the regulation of gene expression. It also promotes inflammation and innate immune responses via NFkB signalling, and may be involved in B cell differentiation and activation^43^. There is some evidence linking increased PARP1 activity to autoimmunity. Specific PARP1 haplotypes are associated with RA^44^. Some, but not all, EAE studies have demonstrated efficacy of PARP1 inhibitors in limiting disease severity^43^. In our study, surprisingly, the direction of SMR effect implies that increased PARP1 expression is protective against MS (Beta_SMR_ −0.31, 2.95×10^−195^).

Data-driven efforts to prioritise drug targets in MS using SMR have been limited to date. One study used older GWAS data and smaller eQTL datasets (Westra *et al* and CAGE) to prioritise 10 non-MHC genes and 20 non-MHC methylation loci associated with MS in peripheral blood ^45^. Our use of larger, more recent datasets, a more lenient HEIDI threshold, and an FDR cutoff rather than a Bonferroni correction is likely to explain our discovery of a greater number of genes. To our knowledge, ours is the first attempt to synthesise SMR with a curated list of druggable targets to prioritise genes for therapy in MS.

An important limitation is that our analyses focus on whole blood, partly because this is the tissue for which the largest eQTL and mQTL datasets exist. As the genetic determinants of gene expression vary between tissues, this paper focuses on identifying differentially-expressed genes in blood potentially suitable for therapeutic interventions. Not only does this miss possible tissue-specific effects of other genes in MS, it also does not distinguish between different types of immune cells. Although MS GWAS hits are primarily enriched for eQTLs and epigenomic marks of active chromatin in various types of immune cells, which are included in eQTL datasets from peripheral blood, there is also enrichment in CNS-resident microglia and, at the tissue level, in the thymus.^1^ Our analysis would not detect cell type-specific effects acting in other tissues (e.g. CNS and thymus), and cell type-specific effects within particular leukocyte subsets which are not reflected in ‘averaged’ datasets across multiple cell types. It is likely that the pathogenesis of MS involves dysregulated signalling in specific subsets of immune cells, and our work misses this complexity. A useful extension of our work would be to apply this approach to eQTL datasets garnered from specific immune cell types.

A further cell type we explore using eQTL data (mQTL data are not available) is LCLs, an *in vitro* model of EBV-infected B cells. This analysis allows us to make inferences about genes which may govern MS susceptibility within EBV-infected B cells. We prioritise 7 genes from this analysis, several of which have biologically plausible mechanisms linking their expression to MS susceptibility. The clearest of these is *TNFRSF14*: a herpes virus entry mediator, which facilitates infection of B lymphocytes by herpes viruses (e.g. EBV and HHV6) and lies within an MS risk locus ^1,46^. The relatively weak correlation between SMR signals in LCLs and whole blood emphasises the diversity of relationships between gene expression and phenotype between different cell types. The importance of MS risk loci acting within EBV-infected cells is reinforced by evidence that MS susceptibility SNPs are enriched for loci influencing EBV copy number and EBV micro-RNA expression within LCLs^47^.

A further limitation of our work is that SMR identifies genes in which the level of expression is associated with disease. There are many ways whereby gene and protein function may influence phenotype independently of levels of gene expression, such as through changes in subcellular localisation and post-translational modifications. In addition, the effect of expression of a single gene is likely to depend on the overall transcriptional and translational state of the cell at that point; this complexity is not captured by considering the expression of individual genes separately. Furthermore, the curated list of druggable genes does not imply that a gene is necessarily a sensible target; some prioritised genes with multiple roles in cell signalling, such as CD40, may be challenging to target in practice without unacceptable off-target effects.

In summary, using data-driven mendelian randomisation and publicly-available datasets, we highlight several possible drug targets for multiple sclerosis. In particular, we highlight a central role for the CD40 signalling pathway, which could be targeted with small molecule compounds already being trialled for other indications.

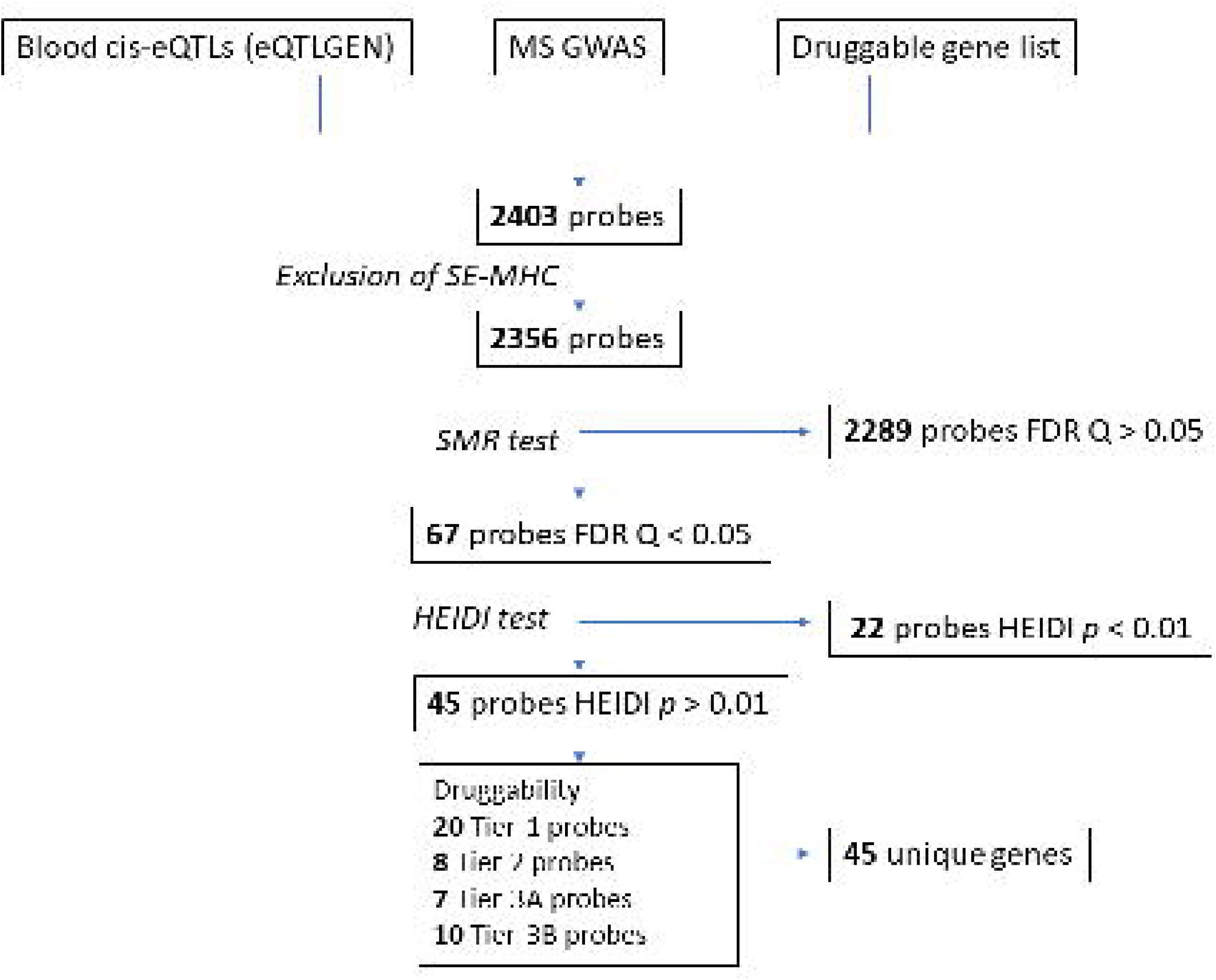

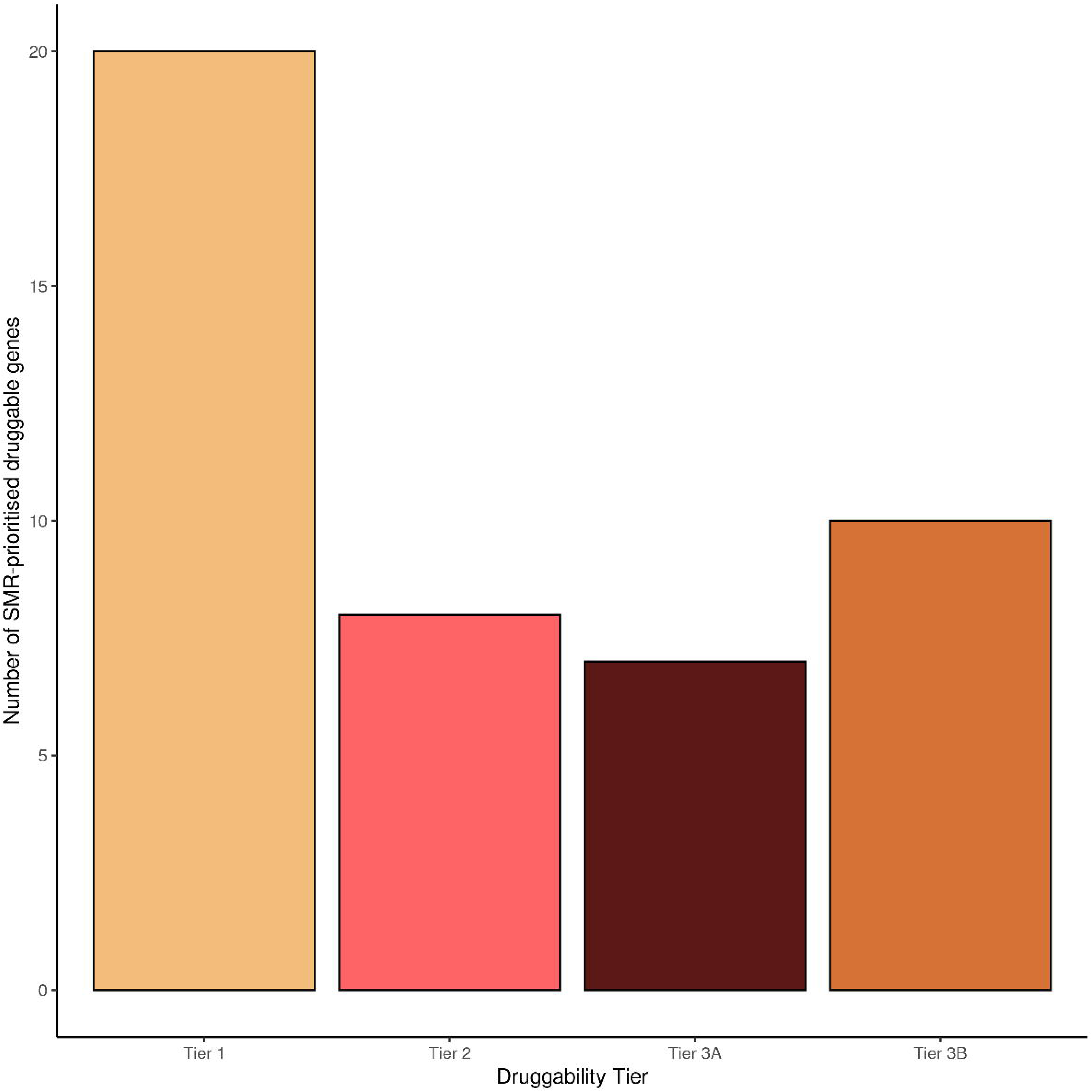

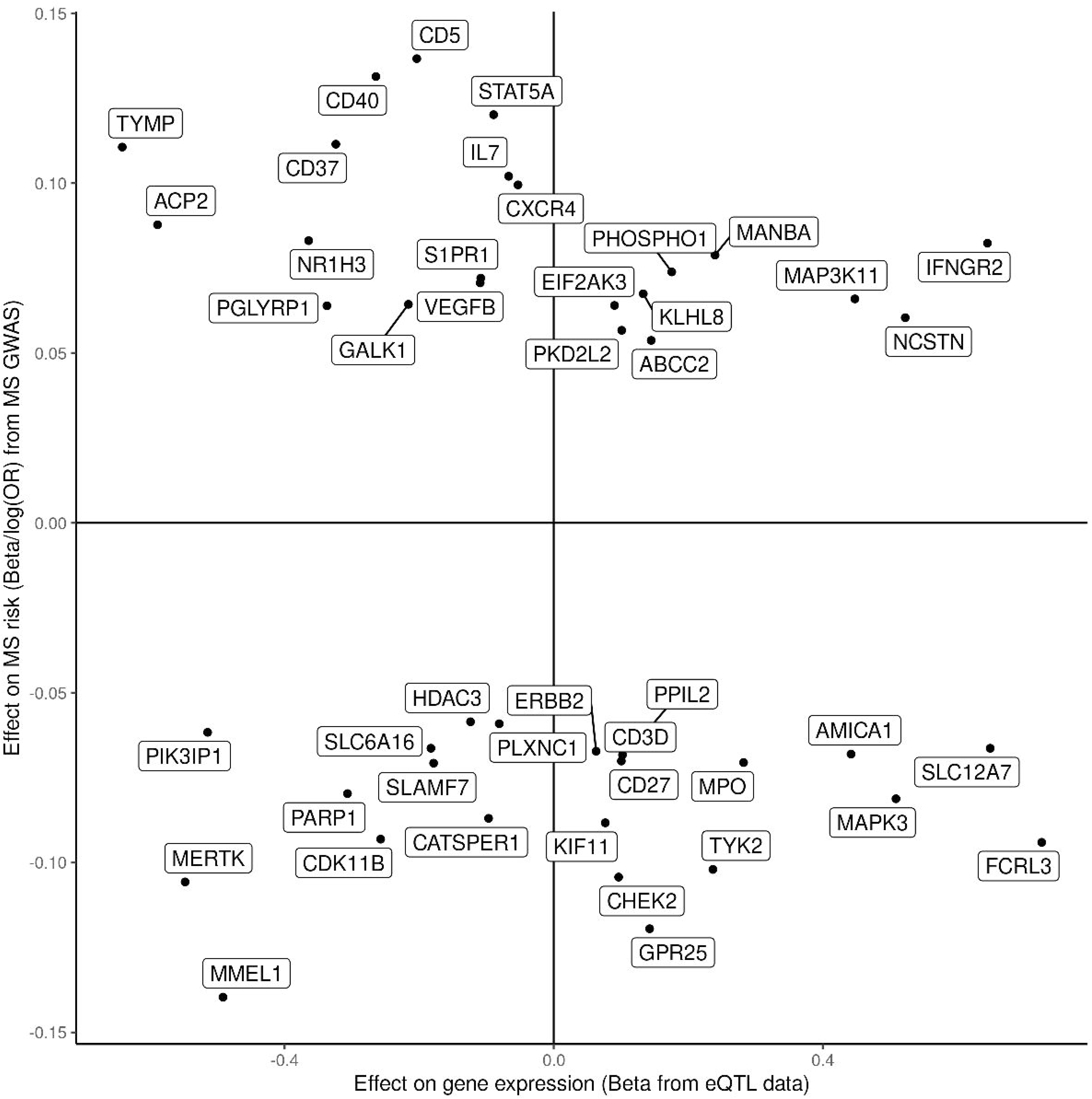

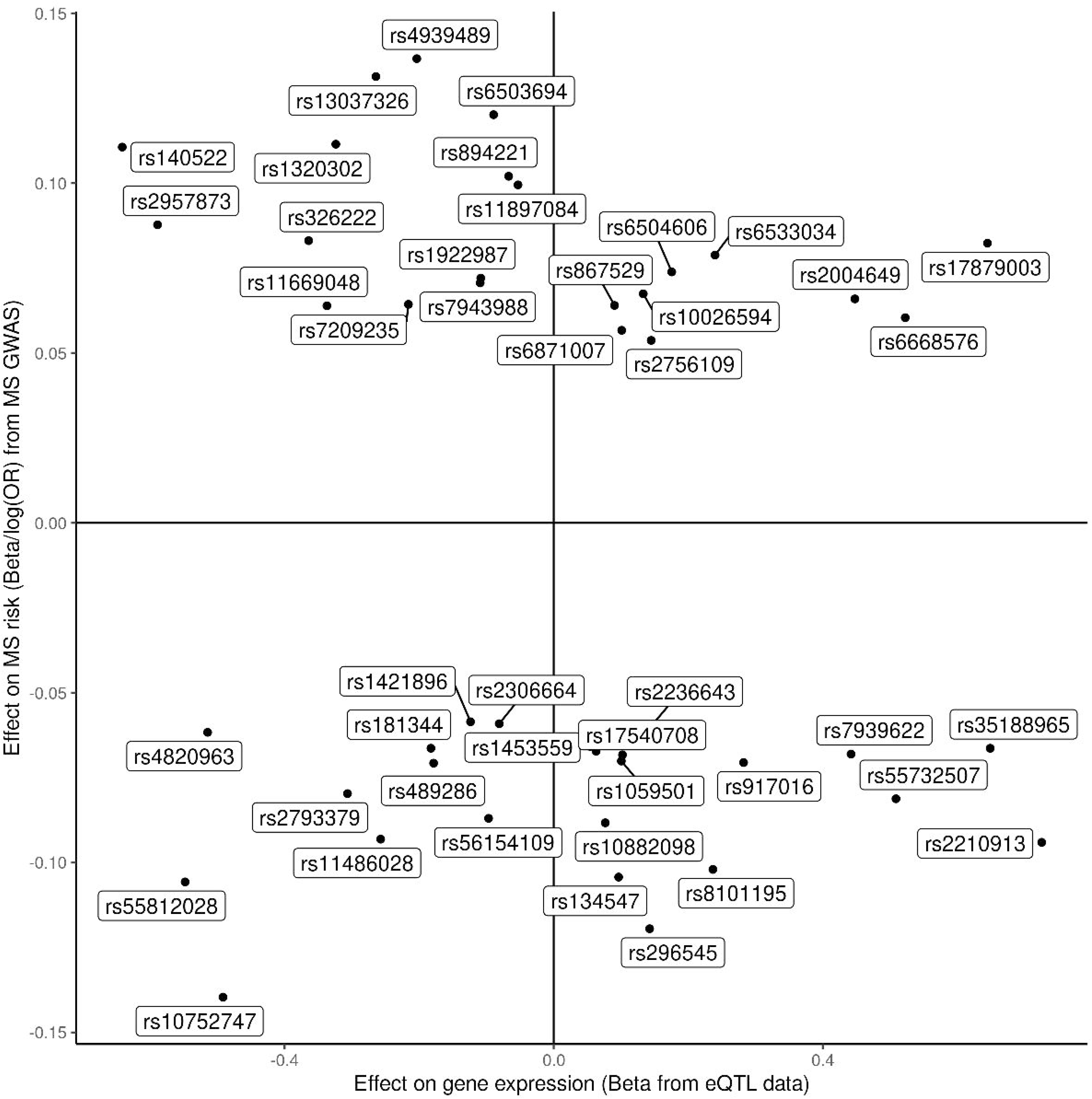

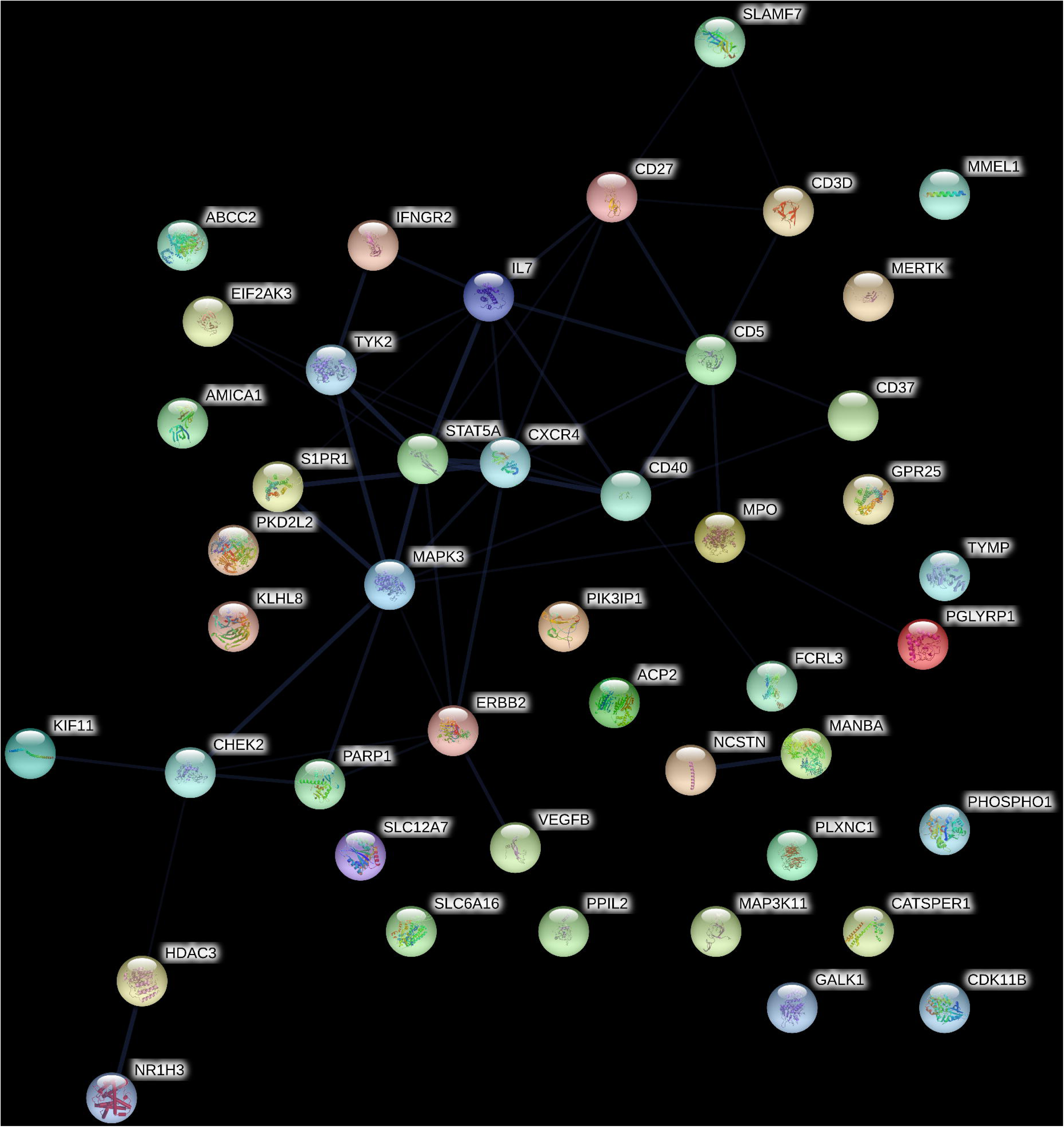

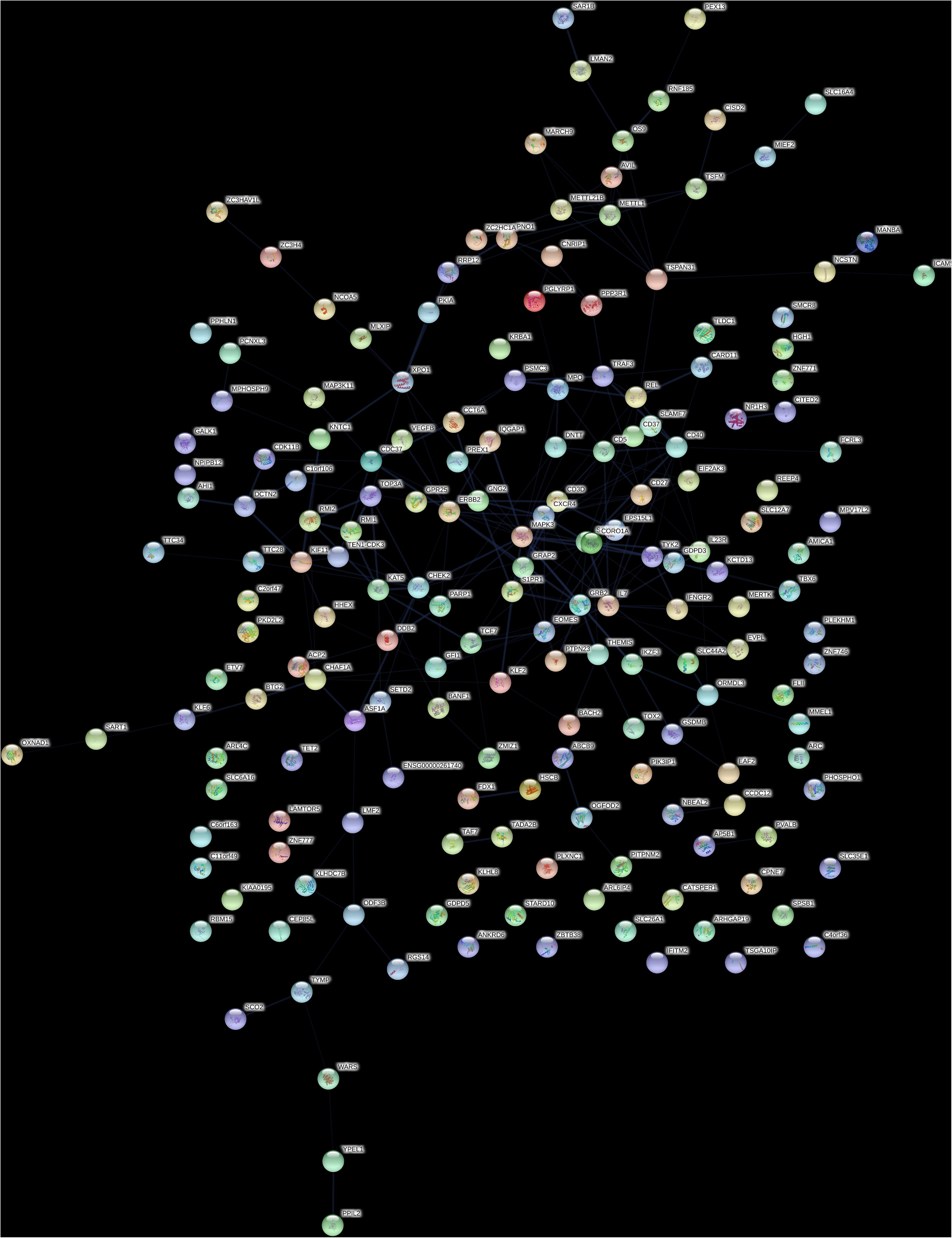

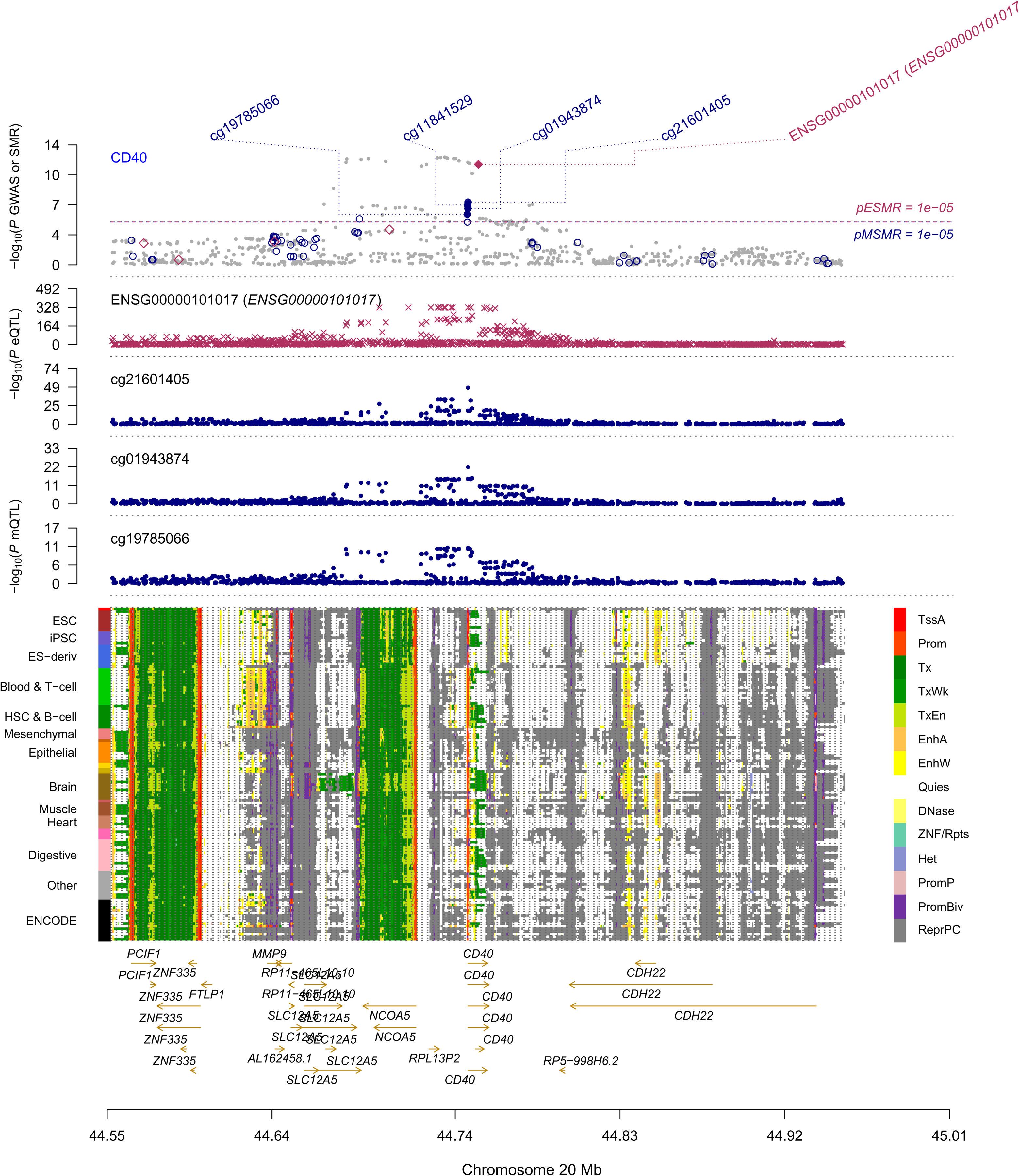

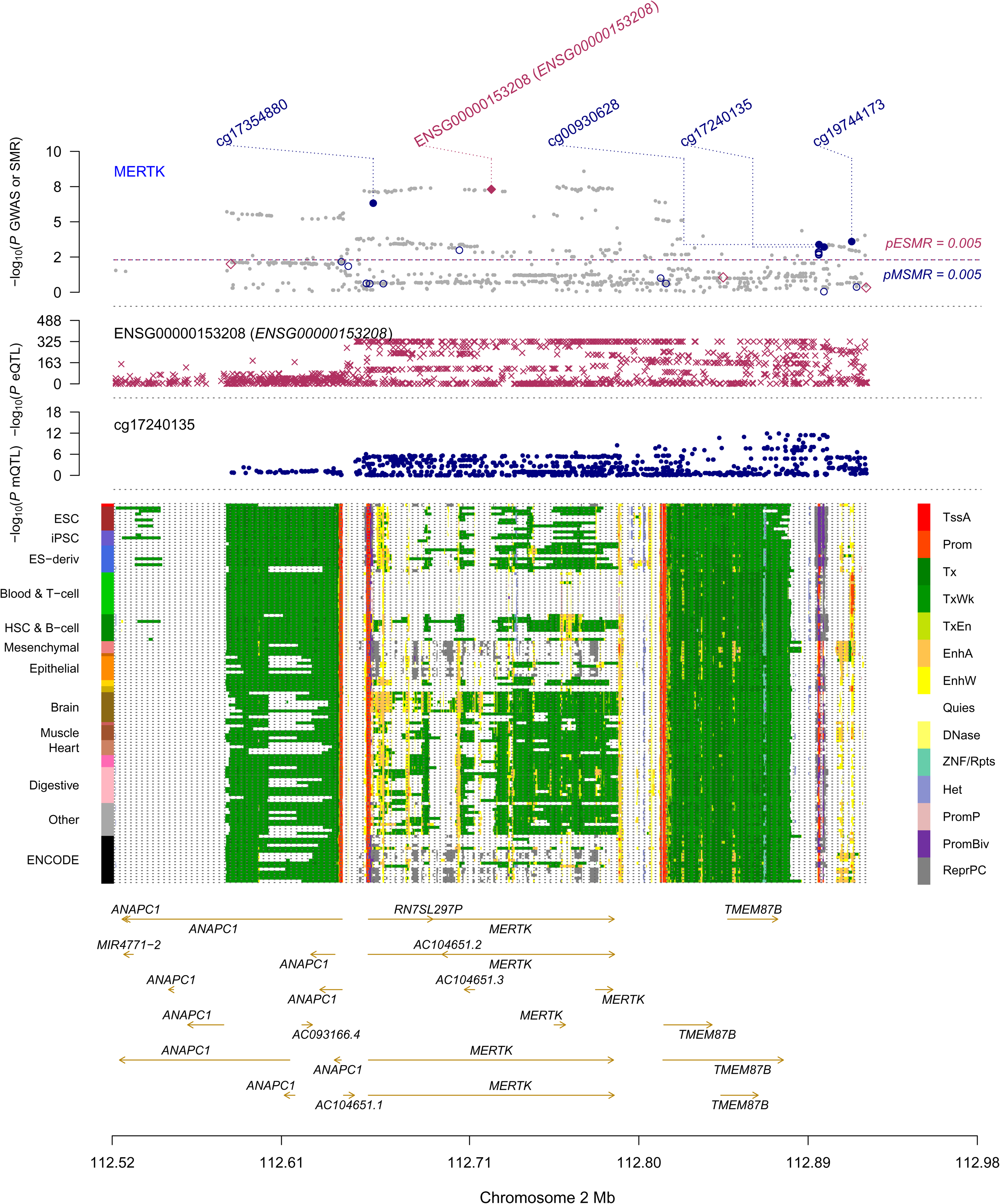

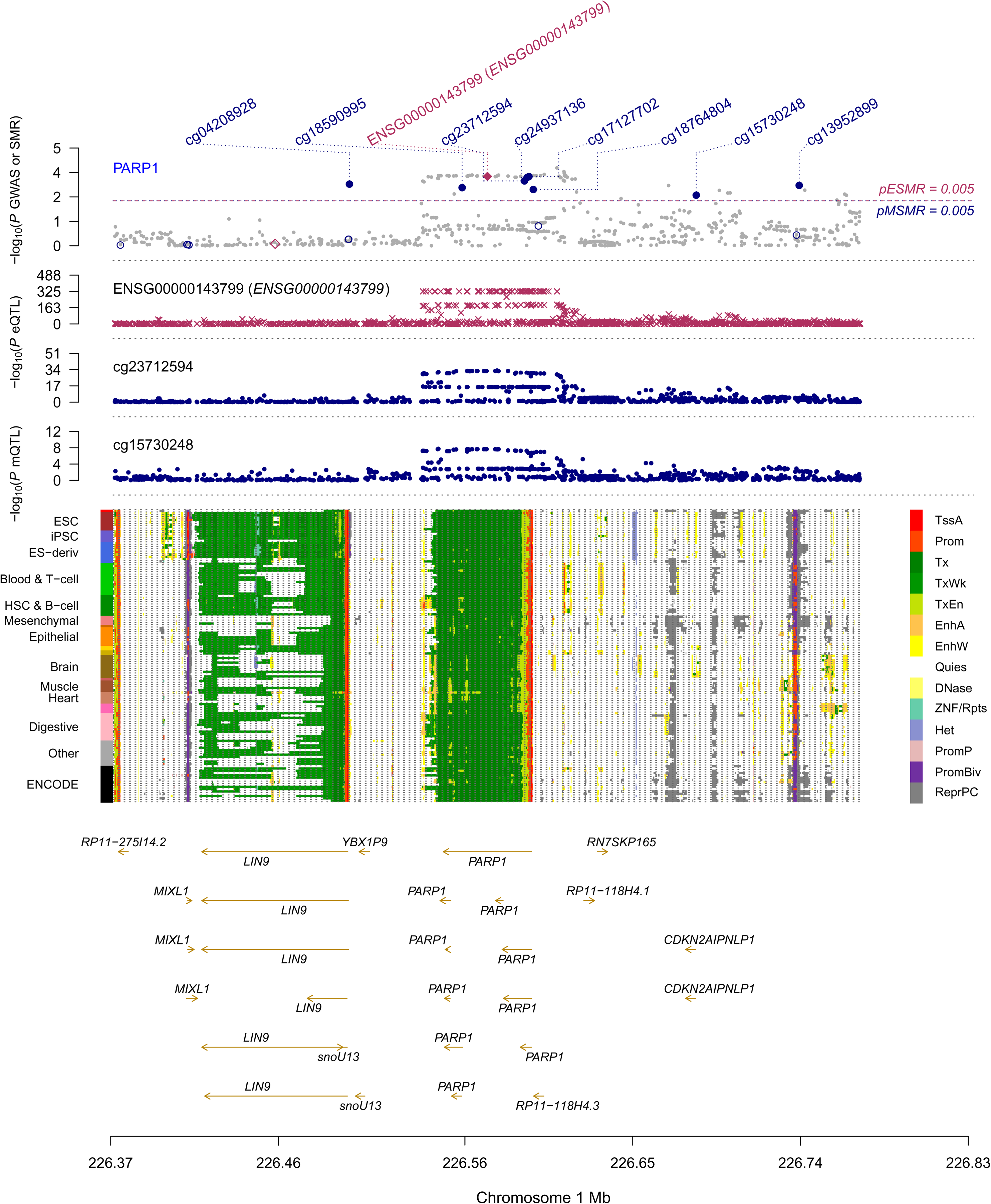

## References

1. International Multiple Sclerosis Genetics Consortium. Multiple sclerosis genomic map implicates peripheral immune cells and microglia in susceptibility. Science 365, (2019).

2. Wu, Y. et al. Integrative analysis of omics summary data reveals putative mechanisms underlying complex traits. Nat. Commun. 9, 918 (2018).

3. Tintore, M., Vidal-Jordana, A. & Sastre-Garriga, J. Treatment of multiple sclerosis - success from bench to bedside. Nat. Rev. Neurol. 15, 53–58 (2019).

4. Zhu, Z. et al. Integration of summary data from GWAS and eQTL studies predicts complex trait gene targets. Nat. Genet. 48, 481–487 (2016).

5. Finan, C. et al. The druggable genome and support for target identification and validation in drug development. Sci. Transl. Med. 9, (2017).

6. Consortium, T. 1000 G. P. & The 1000 Genomes Project Consortium. A global reference for human genetic variation. Nature vol. 526 68–74 (2015).

7. Võsa, U., Claringbould, A., Westra, H. J. & Bonder, M. J. Unraveling the polygenic architecture of complex traits using blood eQTL meta-analysis. bioRxiv (2018).

8. Lloyd-Jones, L. R. et al. The Genetic Architecture of Gene Expression in Peripheral Blood. Am. J. Hum. Genet. 100, 228–237 (2017).

9. Lappalainen, T. et al. Transcriptome and genome sequencing uncovers functional variation in humans. Nature 501, 506–511 (2013).

10. McRae, A. F. et al. Identification of 55,000 Replicated DNA Methylation QTL. Sci. Rep. 8, 17605 (2018).

11. von Mering, C. et al. STRING: known and predicted protein-protein associations, integrated and transferred across organisms. Nucleic Acids Res. 33, D433–7 (2005).

12. Szklarczyk, D. et al. STRING v11: protein-protein association networks with increased coverage, supporting functional discovery in genome-wide experimental datasets. Nucleic Acids Res. 47, D607–D613 (2019).

13. Franceschini, A. et al. STRING v9.1: protein-protein interaction networks, with increased coverage and integration. Nucleic Acids Res. 41, D808–15 (2013).

14. Rivals, I., Personnaz, L., Taing, L. & Potier, M.-C. Enrichment or depletion of a GO category within a class of genes: which test? Bioinformatics 23, 401–407 (2007).

15. Li, T. et al. GeNets: a unified web platform for network-based genomic analyses. Nat. Methods 15, 543–546 (2018).

16. King, T., Butcher, S. & Zalewski, L. Apocrita - High Performance Computing Cluster for Queen Mary University of London. (2017). doi:10.5281/zenodo.438045.

17. Kappos, L. et al. A placebo-controlled trial of oral fingolimod in relapsing multiple sclerosis. N. Engl. J. Med. 362, 387–401 (2010).

18. Cohen, J. A. et al. Oral fingolimod or intramuscular interferon for relapsing multiple sclerosis. N. Engl. J. Med. 362, 402–415 (2010).

19. Calabresi, P. A. et al. Safety and efficacy of fingolimod in patients with relapsing-remitting multiple sclerosis (FREEDOMS II): a double-blind, randomised, placebo-controlled, phase 3 trial. Lancet Neurol. 13, 545–556 (2014).

20. Kappos, L. et al. Siponimod versus placebo in secondary progressive multiple sclerosis (EXPAND): a double-blind, randomised, phase 3 study. Lancet 391, 1263–1273 (2018).

21. van Vollenhoven, R. F. et al. Tofacitinib or adalimumab versus placebo in rheumatoid arthritis. N. Engl. J. Med. 367, 508–519 (2012).

22. Gladman, D. et al. Tofacitinib for Psoriatic Arthritis in Patients with an Inadequate Response to TNF Inhibitors. N. Engl. J. Med. 377, 1525–1536 (2017).

23. Sandborn, W. J. et al. Tofacitinib as Induction and Maintenance Therapy for Ulcerative Colitis. N. Engl. J. Med. 376, 1723–1736 (2017).

24. Aarts, S. A. B. M. et al. The CD40-CD40L Dyad in Experimental Autoimmune Encephalomyelitis and Multiple Sclerosis. Front. Immunol. 8, 1791 (2017).

25. Field, J. et al. The MS Risk Allele of CD40 Is Associated with Reduced Cell-Membrane Bound Expression in Antigen Presenting Cells: Implications for Gene Function. PLoS One 10, e0127080 (2015).

26. Smets, I. et al. Multiple sclerosis risk variants alter expression of co-stimulatory genes in B cells. Brain 141, 786–796 (2018).

27. Boumpas, D. T. et al. A short course of BG9588 (anti-CD40 ligand antibody) improves serologic activity and decreases hematuria in patients with proliferative lupus glomerulonephritis. Arthritis Rheum. 48, 719–727 (2003).

28. André, P. et al. CD40L stabilizes arterial thrombi by a beta3 integrin--dependent mechanism. Nat. Med. 8, 247–252 (2002).

29. Robles-Carrillo, L. et al. Anti-CD40L immune complexes potently activate platelets in vitro and cause thrombosis in FCGR2A transgenic mice. J. Immunol. 185, 1577–1583 (2010).

30. Tocoian, A. et al. First-in-human trial of the safety, pharmacokinetics and immunogenicity of a PEGylated anti-CD40L antibody fragment (CDP7657) in healthy individuals and patients with systemic lupus erythematosus. Lupus 24, 1045–1056 (2015).

31. Karnell, J. L. et al. A CD40L-targeting protein reduces autoantibodies and improves disease activity in patients with autoimmunity. Sci. Transl. Med. 11, (2019).

32. Michel, N. A., Zirlik, A. & Wolf, D. CD40L and Its Receptors in Atherothrombosis-An Update. Front Cardiovasc Med 4, 40 (2017).

33. Kotelnikova, E. et al. MAPK pathway and B cells overactivation in multiple sclerosis revealed by phosphoproteomics and genomic analysis. Proc. Natl. Acad. Sci. U. S. A. 116, 9671–9676 (2019).

34. Arthur, J. S. C. & Ley, S. C. Mitogen-activated protein kinases in innate immunity. Nat. Rev. Immunol. 13, 679–692 (2013).

35. Wium, M., Paccez, J. D. & Zerbini, L. F. The Dual Role of TAM Receptors in Autoimmune Diseases and Cancer: An Overview. Cells 7, (2018).

36. Scott, R. S. et al. Phagocytosis and clearance of apoptotic cells is mediated by MER. Nature 411, 207–211 (2001).

37. Chung, W.-S. et al. Astrocytes mediate synapse elimination through MEGF10 and MERTK pathways. Nature 504, 394–400 (2013).

38. Binder, M. D. et al. Gas6 deficiency increases oligodendrocyte loss and microglial activation in response to cuprizone-induced demyelination. J. Neurosci. 28, 5195–5206 (2008).

39. Binder, M. D. et al. Gas6 increases myelination by oligodendrocytes and its deficiency delays recovery following cuprizone-induced demyelination. PLoS One 6, e17727 (2011).

40. Weinger, J. G., Omari, K. M., Marsden, K., Raine, C. S. & Shafit-Zagardo, B. Up-regulation of soluble Axl and Mer receptor tyrosine kinases negatively correlates with Gas6 in established multiple sclerosis lesions. Am. J. Pathol. 175, 283–293 (2009).

41. Sainaghi, P. P. et al. Growth arrest specific gene 6 protein concentration in cerebrospinal fluid correlates with relapse severity in multiple sclerosis. Mediators Inflamm. 2013, 406483 (2013).

42. Lee-Sherick, A. B. et al. MERTK inhibition alters the PD-1 axis and promotes anti-leukemia immunity. JCI Insight 3, (2018).

43. Rosado, M. M., Bennici, E., Novelli, F. & Pioli, C. Beyond DNA repair, the immunological role of PARP-1 and its siblings. Immunology 139, 428–437 (2013).

44. Pascual, M. et al. A poly(ADP-ribose) polymerase haplotype spanning the promoter region confers susceptibility to rheumatoid arthritis: PARP-1 Gene Promoter Polymorphism and RA Susceptibility. Arthritis & Rheumatism 48, 638–641 (2003).

45. Mo, X.-B. et al. Integrative analysis revealed potential causal genetic and epigenetic factors for multiple sclerosis. J. Neurol. 266, 2699–2709 (2019).

46. Blanco-Kelly, F. et al. Members 6B and 14 of the TNF receptor superfamily in multiple sclerosis predisposition. Genes Immun. 12, 145–148 (2011).

47. Afrasiabi, A., Parnell, G. P., Swaminathan, S., Stewart, G. J. & Booth, D. R. The interaction of Multiple Sclerosis risk loci with Epstein-Barr virus phenotypes implicates the virus in pathogenesis. Sci. Rep. 10, 193 (2020).

