## Supplementary figures and images for "Summary-data-based mendelian randomisation reveals druggable targets for multiple sclerosis"

### supplementary_figure1.png

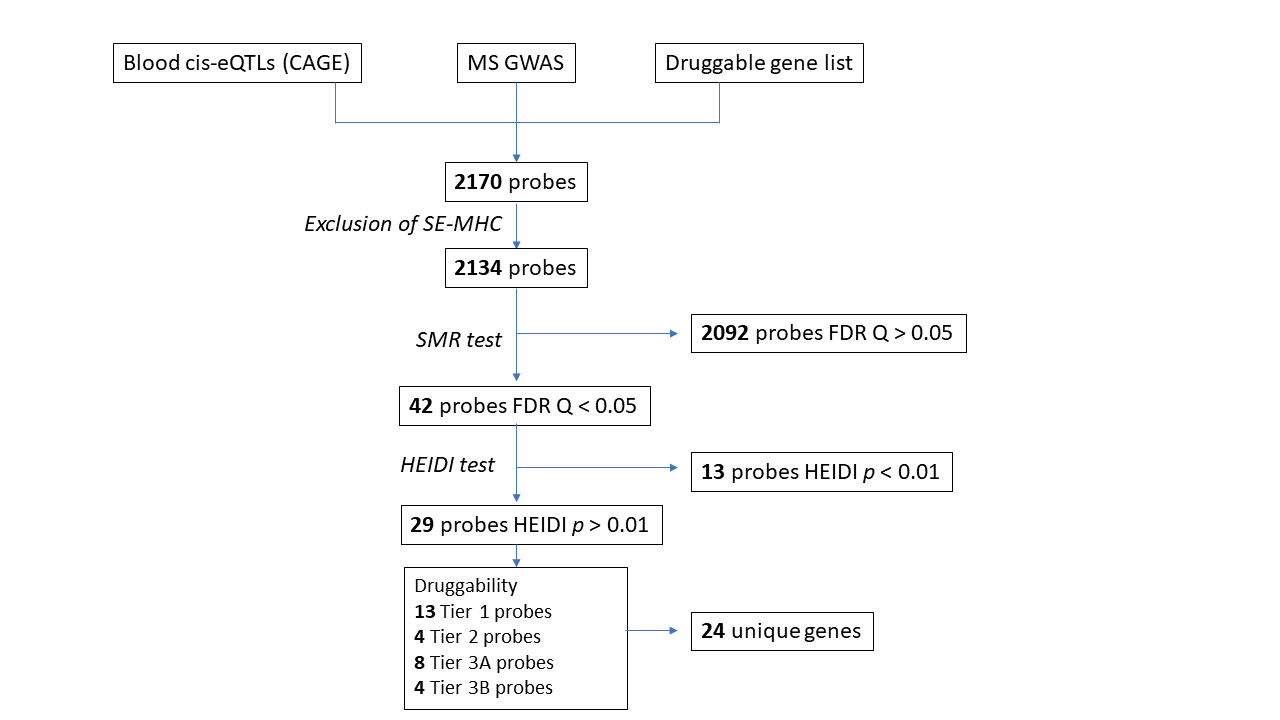

### supplementary_figure3.png

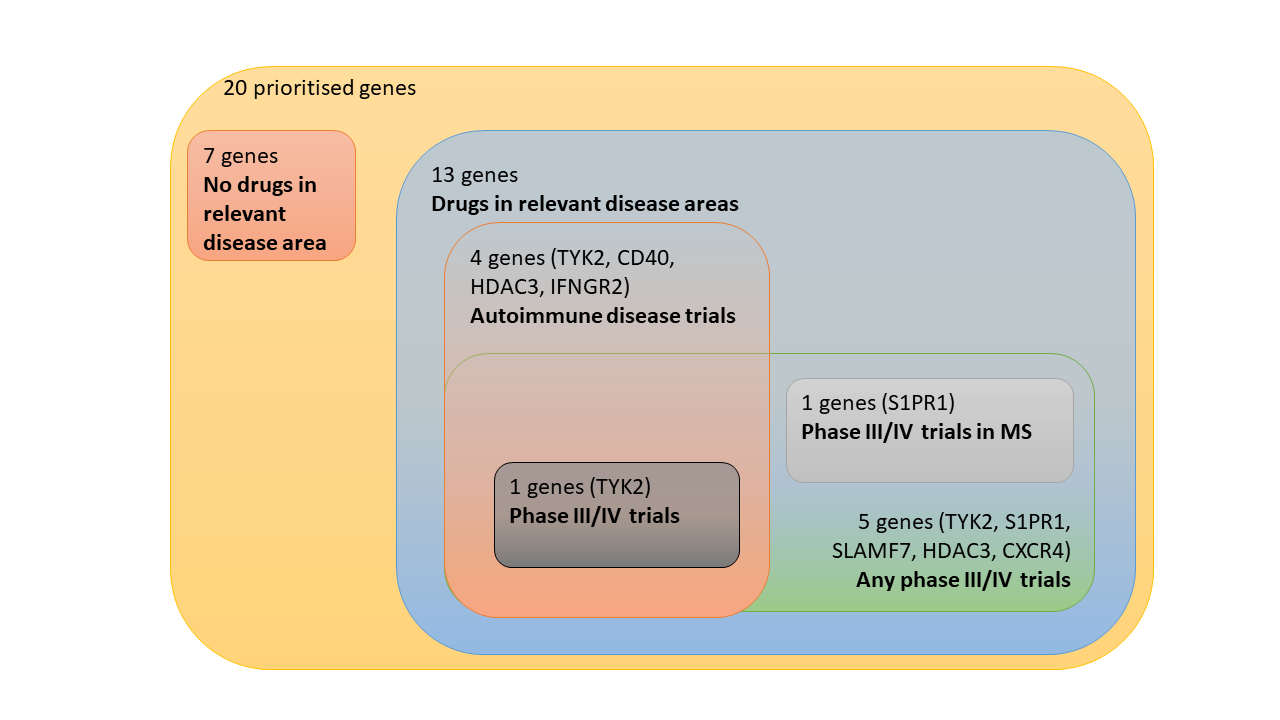

### supplementary_figure4.png

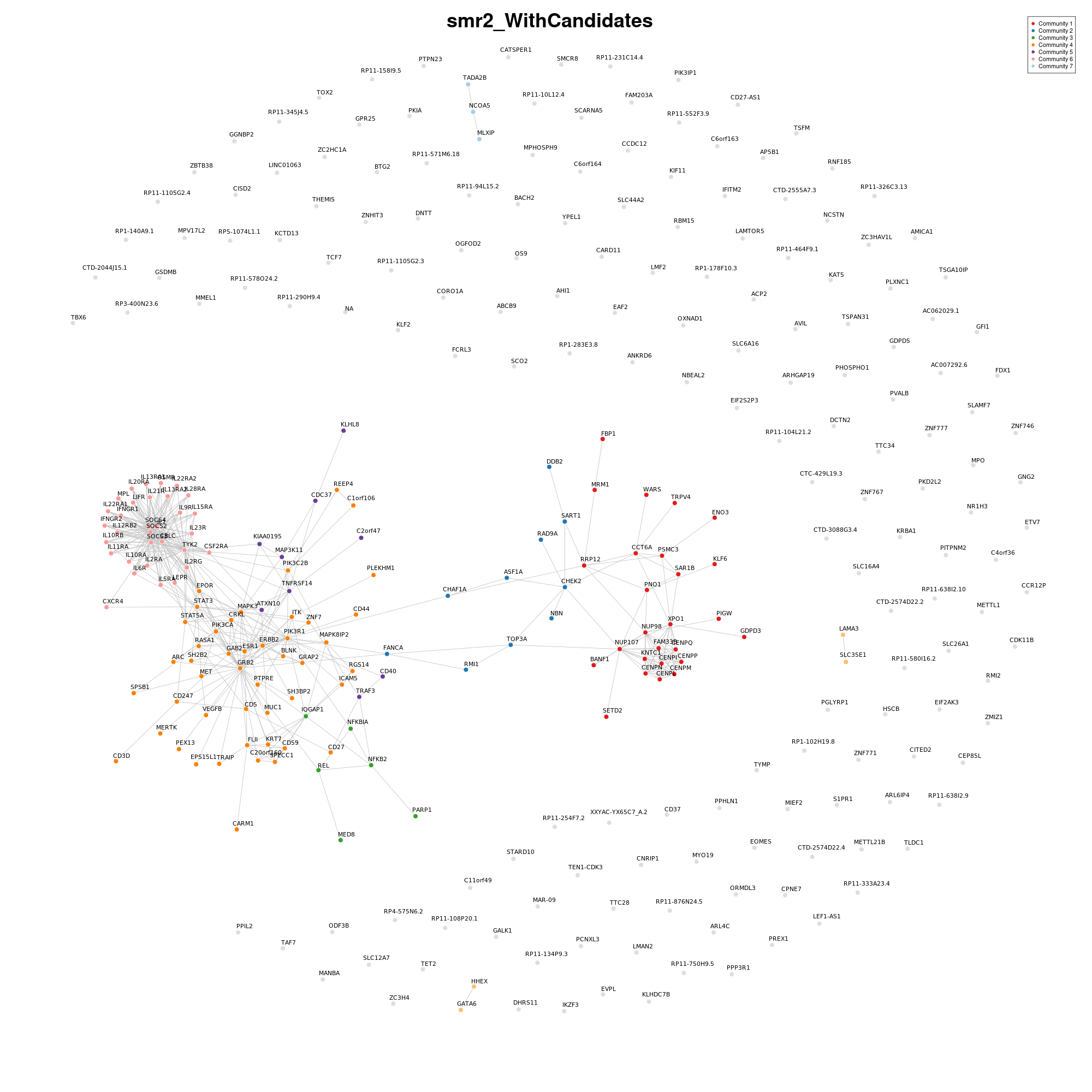

### supplementary_figure5a.png

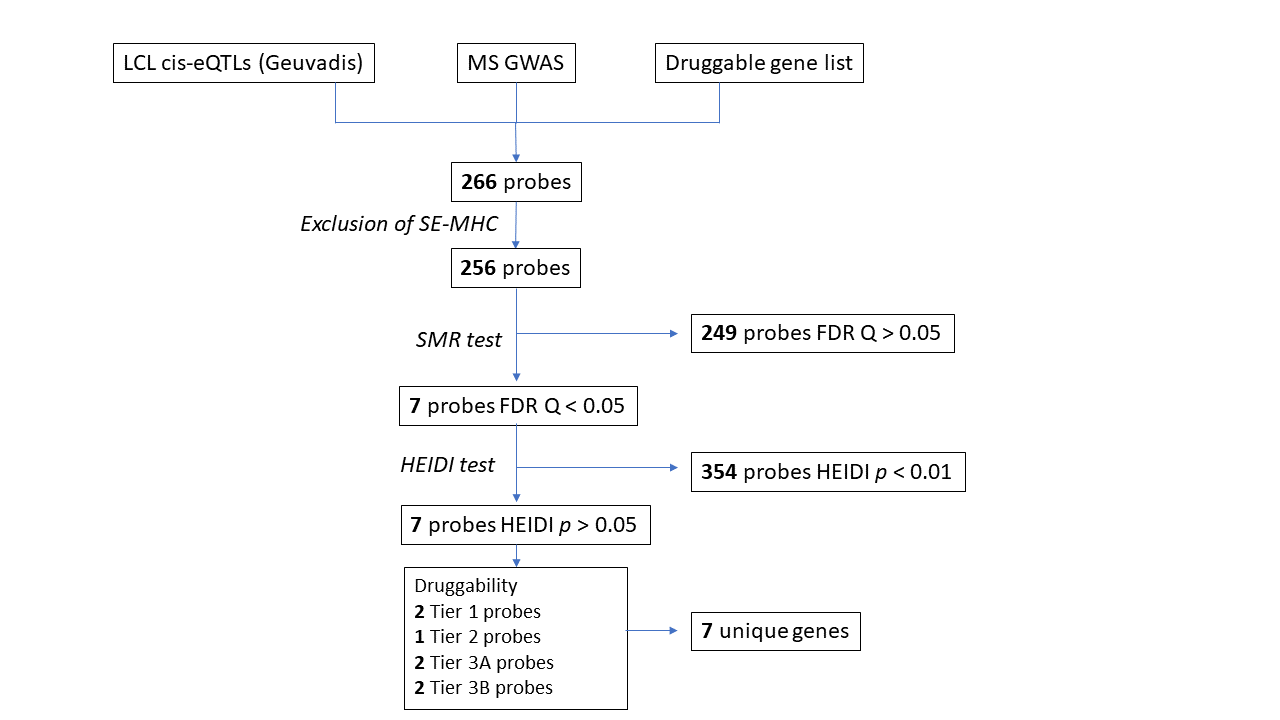

### supplementary_figure5b.png

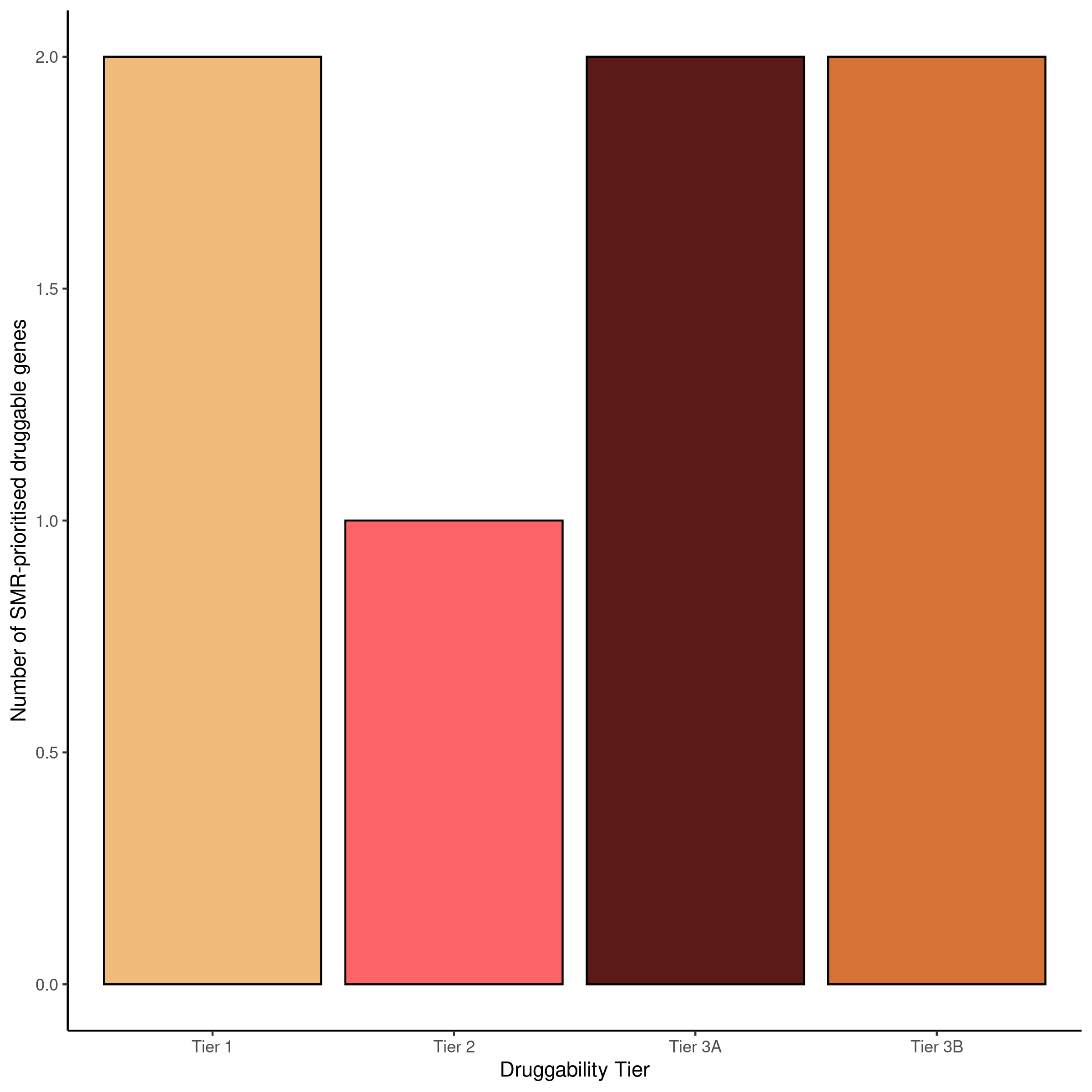

### supplementary_figure5c.png

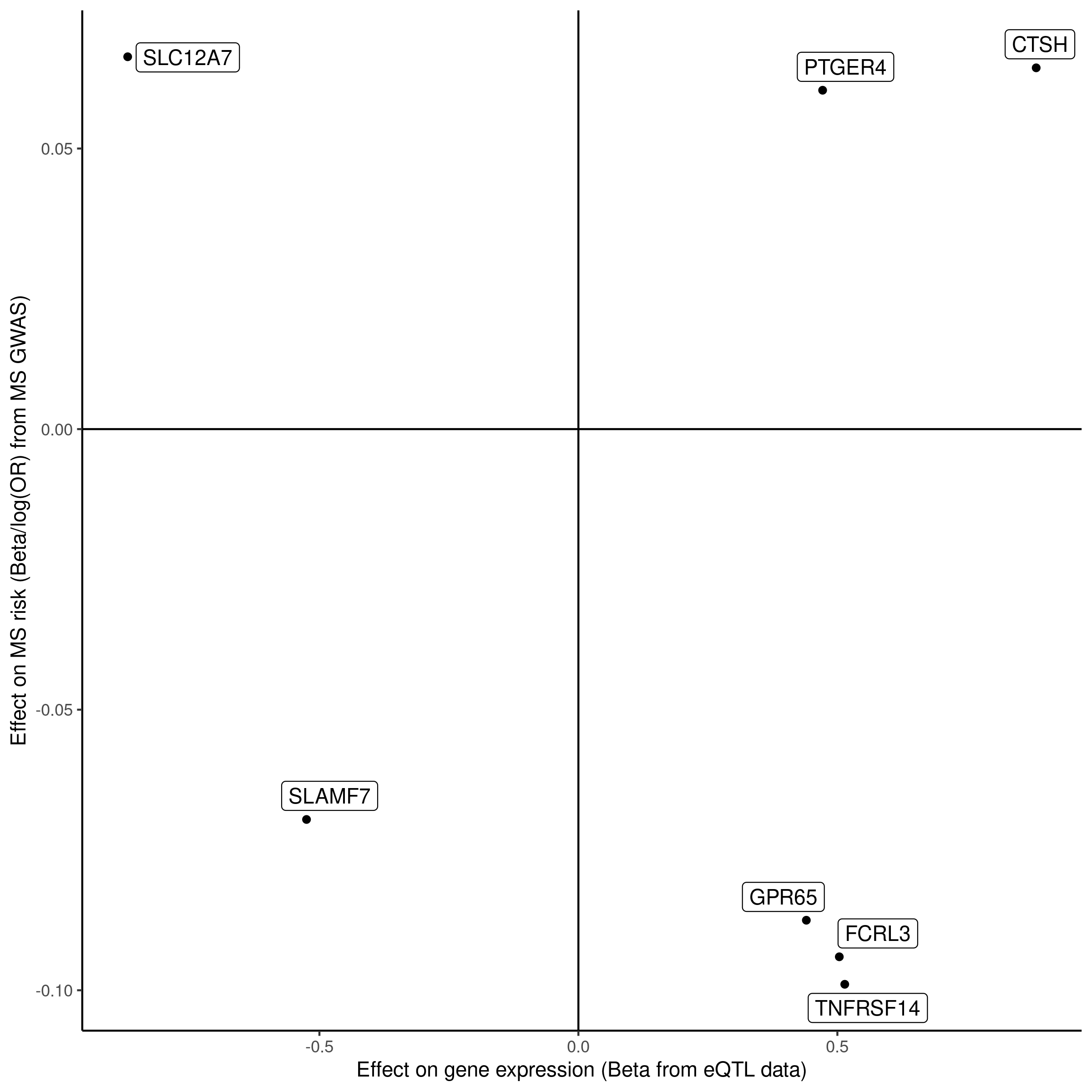

### supplementary_figure5d.png

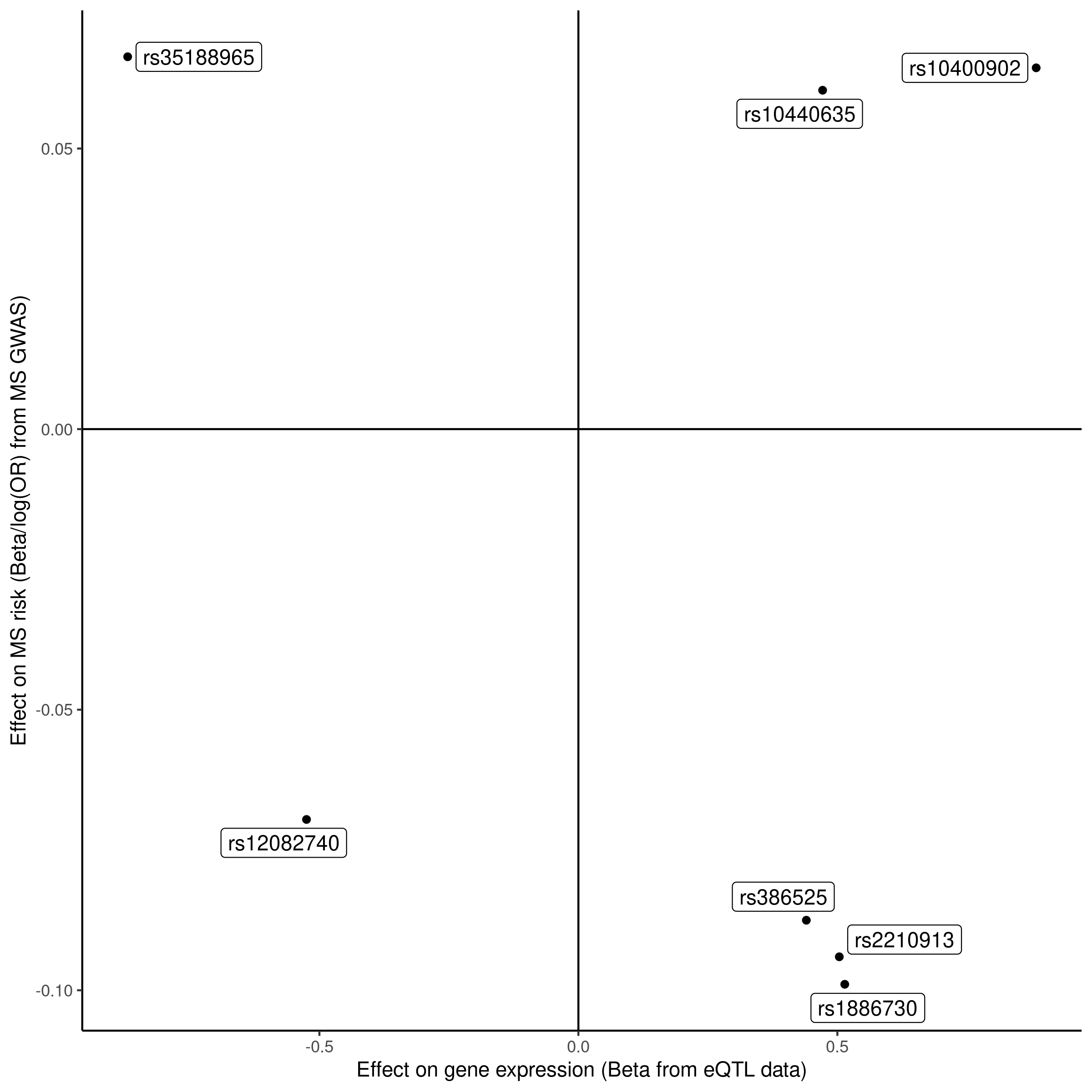

### supplementary_figure_2a.png

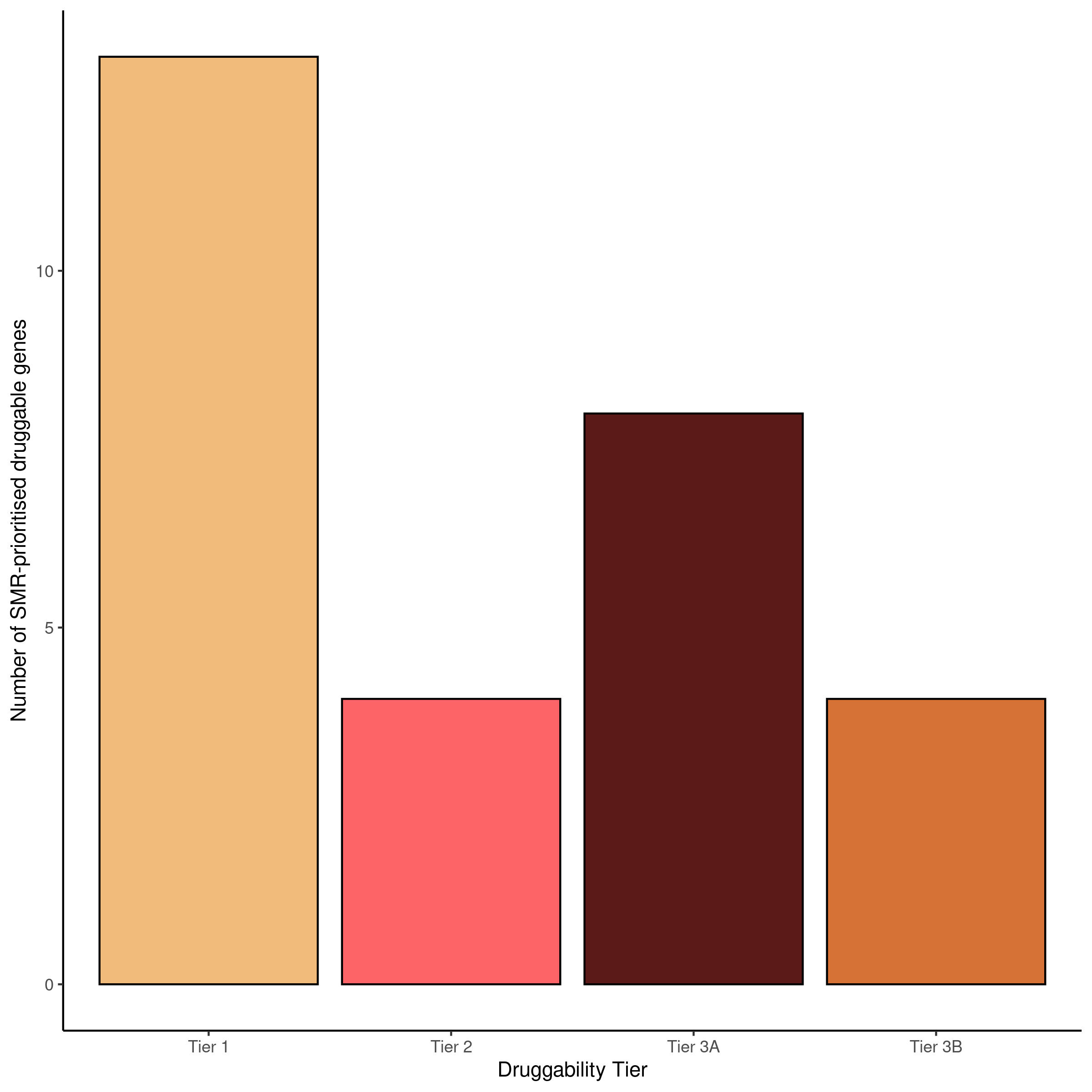

### supplementary_figure_2b.png

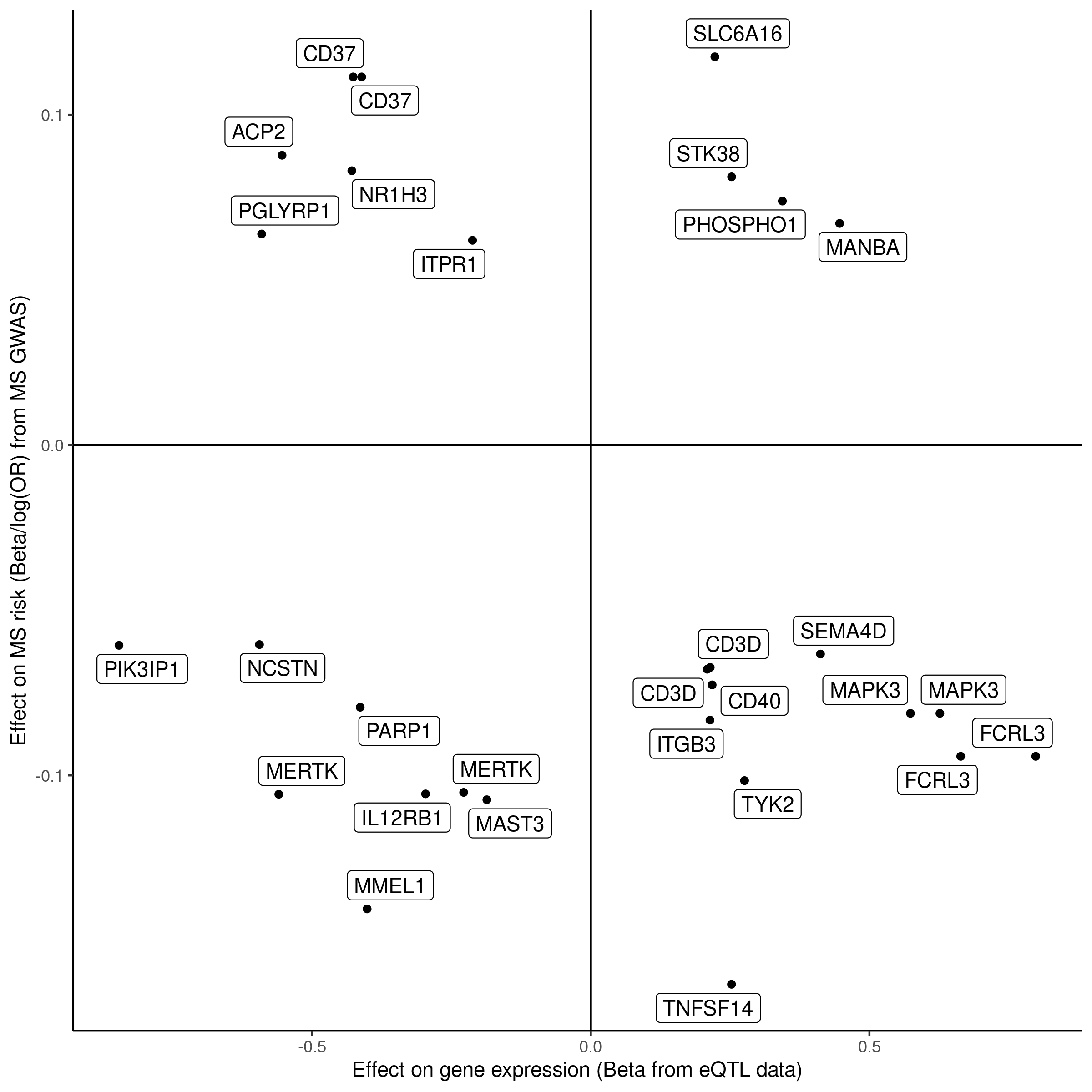

### supplementary_figure_2c.png

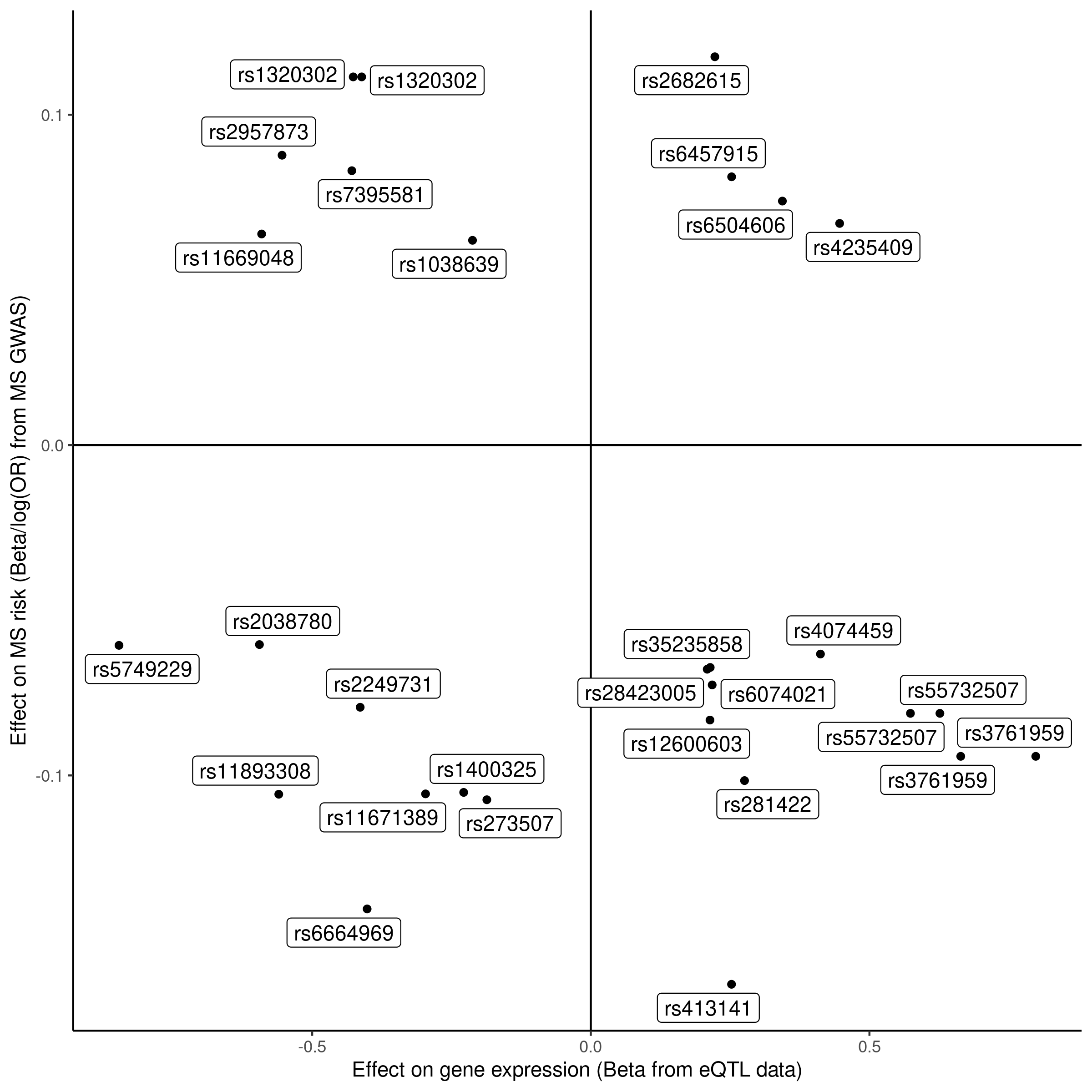

### supplementary_figure_2d.png

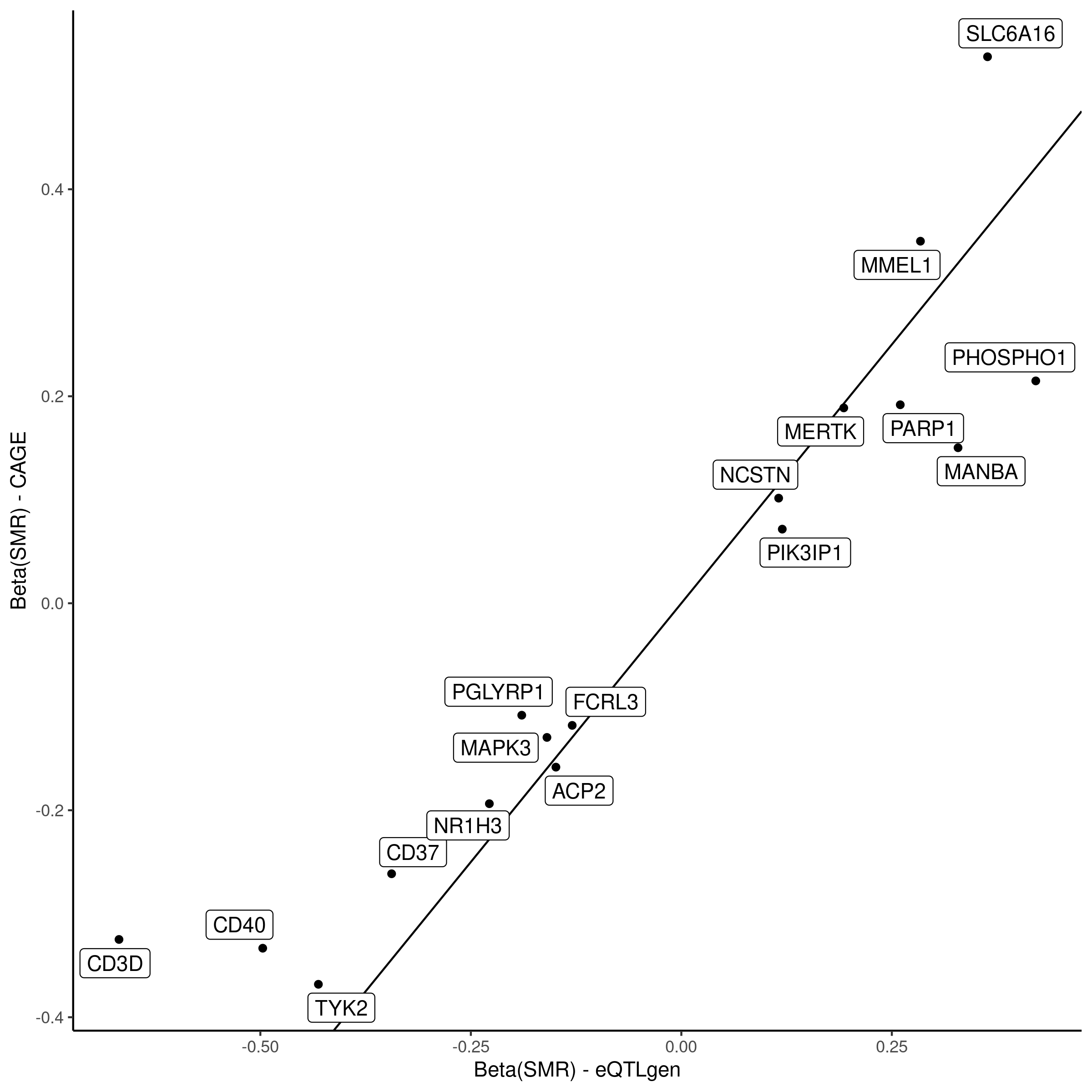
